# Focal adhesions are controlled by microtubules through local contractility regulation

**DOI:** 10.1101/2023.04.17.535593

**Authors:** J. Aureille, S.F.H. Barnett, I. Arnal, L. Lafanechère, B.C. Low, P. Kanchanawong, A. Mogilner, A.D. Bershadsky

## Abstract

Microtubules regulate cell polarity and migration by local activation of focal adhesion turnover, but the mechanism of this process is insufficiently understood. Molecular complexes containing KANK family proteins connect microtubules with the major component of focal adhesions, talin. Local optogenetic activation of KANK1-mediated links which promoted microtubule targeting to individual focal adhesion resulting in its centripetal sliding and rapid disassembly. The sliding is preceded by a local increase of traction force due to accumulation of myosin-II and actin in the proximity of the focal adhesion. Knockdown of Rho activator GEF-H1 prevented development of traction force and abolished sliding and disassembly of focal adhesion upon KANK activation. Other players participating in microtubule-driven KANK-dependent focal adhesion disassembly include kinases ROCK and PAK, as well as microtubules/focal adhesions associated proteins Kinesin-1, APC and αTAT. Finally, we propose a physical model of a microtubule-driven focal adhesion disruption involving local GEF-H1/RhoA/ROCK dependent activation of contractility which is consistent with experimental data.

## Introduction

Microtubules are ubiquitous cytoskeletal elements. In many cell types, they are responsible for polarity of cell shape and cell directional migration^1–3^. This function of microtubules depends on their interaction with integrin-mediated cell-matrix adhesions. In this study, we focus on microtubule interaction with a special type of integrin adhesions known as focal adhesions.

Focal adhesions are associated with the contractile actomyosin cytoskeleton and transduce forces generated by associated actomyosin structures (stress fibers) to the extracellular matrix. A variety of experimental approaches based on the use of elastic deformable substrate (traction force microscopy) clearly demonstrated that many cell types exert forces applied to the substate at discrete points corresponding to focal adhesions^4–6^. At the same time, the focal adhesions are also sensing the forces: they disassemble when myosin-IIA driven actomyosin contractility is inhibited and grow in size when the contractility increases or external force is applied^6–11^.

Disruption of microtubules results in increase of focal adhesion size in a myosin-II dependent fashion^12^. The mechanism underlying this phenomenon is based on activation of Rho and Rho kinase (ROCK) upon microtubule disruption^13^. This Rho activation occurs due to release of a microtubule-associated Rho GEF, GEF-H1 and its activation^14, 15^. Indeed, GEF-H1 knockdown abolished the growth of focal adhesions upon microtubule disruption^14, 16^.

While the growth and shrinking dynamics of microtubules is faster than turnover of focal adhesions, microtubules remain dynamically linked to focal adhesions through a complex protein network. One of the main players in microtubule coupling to focal adhesions are the KANK family of proteins, which bind to a major focal adhesion component, talin, via their N-terminal KN domains^17, 18^. At the same time, the central coiled coil domains of KANK proteins bind a liprin-β1, a component of membrane-associated complex that include also liprin-α, ELKS and LL5β^1^. LL5β in turn binds the microtubule end tracking proteins CLASP1/2. In addition, ankyrin repeat domain at the C-terminus of KANK binds kif21a kinesin, which is also located at the microtubule plus end^18, 19^. Thus, this complex can trap microtubule ends and link them via KANK to the focal adhesions^20, 21^.

We have shown previously that disruption of the coupling between microtubules and focal adhesions by either knockdown of KANK proteins or by overexpression of KN domain which interferes with the binding of endogenous KANK to talin reproduces the effect of total disruption of microtubules^16^. The GEF-H1 released from microtubules apparently undergoes activation resulting in formation of numerous myosin-II filaments. This in turn, led to increase of focal adhesion sizes similarly to that observed upon total microtubule disruption^16^. These data are consistent with previously published results showing that depletion of other elements in the microtubule-focal adhesion link such as CLASPs^22^ or EB1^23^ as well as ELKS (ERC1) ^24^, results in suppression of focal adhesion turnover. Thus, both microtubule disruption and their disconnection from focal adhesions induce focal adhesion growth in GEF-H1 and myosin-II dependent manner^25^.

In agreement with these results, the microtubule outgrowth after washing out a microtubule disrupting drug leads to transient disassembly of the focal adhesions^25^. Using this experimental model, several candidate proteins related to the microtubule mediated disassembly of focal adhesions were proposed. However, this approach only allowed assessment of the alteration of focal adhesion areas in the entire cell rather than the effect of individual microtubules targeting individual focal adhesions. In addition, since the process of microtubule outgrowth is not entirely synchronous, the time-course of the microtubule driven focal adhesion disassembly was difficult to follow. Finally, these studies did not assess the changes in myosin-II contractility that, as we mention above, are critically important for the understanding of microtubule interactions with focal adhesions.

In the present study we developed an optogenetic approach which permitted us to target microtubules to selected individual focal adhesions. This approach is based on using the iLID optogenetic system^26^ which, upon blue light illumination, restored the link between two halves of KANK molecule promoting the KANK mediated contacts between microtubules and illuminated adhesions. We have shown that microtubule targeting indeed results in focal adhesion disassembly and analyzed this process in detail.

We have found that the process of microtubule driven focal adhesion disassembly requires GEF-H1 dependent local activation of myosin-II filament formation. The burst of actomyosin contractility triggers the disruption of the adhesion by inducing its detachment and sliding. Since the focal adhesion does not experience stretching force during sliding, it undergoes rapid disassembly. We have identified the microtubule- and focal adhesion-associated proteins involved in this process and showed that some of them mediate the burst of contractility triggering the focal adhesion sliding while others likely weakened the adhesion strength making the sliding possible. We propose a plausible quantitative model of the entire process of interactions of microtubules with focal adhesion leading to focal adhesion disassembly.

## Results

### Disassembly of focal adhesions upon induction of KANK-mediated link

We created an iLID system-based optogenetic construct^26^ of KANK1 protein (OptoKANK) which consists of i) KANK1 talin binding domain fused with mApple at its N-terminus and LOV2ssrA at the C-terminus and ii) remaining part of KANK1 molecule (ΔKN) fused with SSpB at the N-terminus and mEmerald at the C-terminus (**figure 1A**). Total Internal reflection fluorescence (TIRF) microscopy of distribution of these constructs in HT1080 cells together with focal adhesion marker vinculin-mIFP revealed that KN domain localized to focal adhesions while ΔKN did not and demonstrated weak surface fluorescence (**figure 1B**). Illumination of a small circular area containing a focal adhesion did not affect localization of KN but led to the relocation of ΔKN into illuminated region in proximity of the focal adhesion (**figure 1B**).

**Figure 1:**
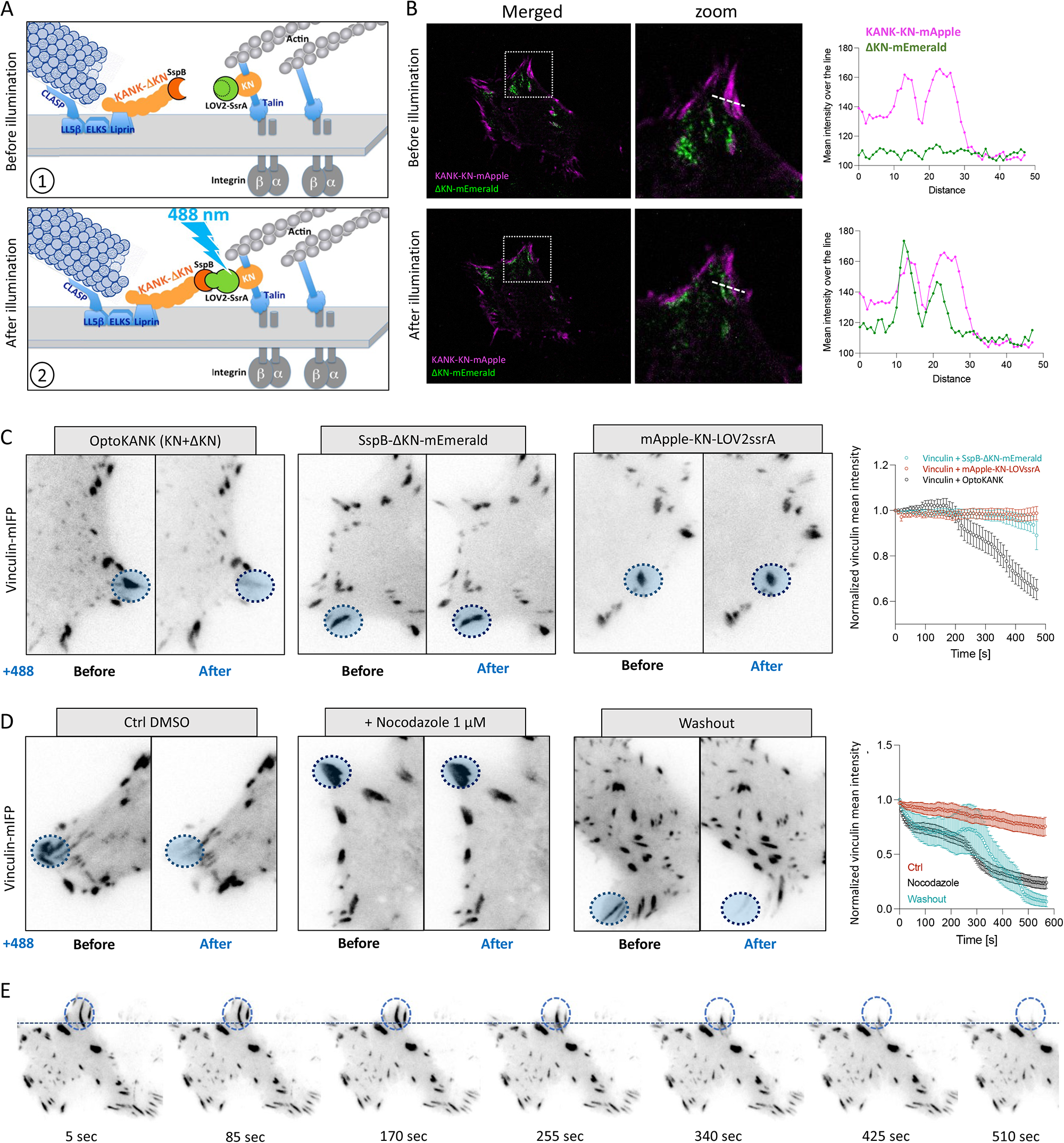
Disassembly of focal adhesions upon induction of KANK-mediated link. **(A)** Schematic illustration of OptoKANK activation. 1) In dark state the C-terminal helix of the LOV2 domain prevents the SsrA peptide to bind its binding partner SspB. 2) Upon blue illumination (488nm) LOV2 changes its conformation allowing SsrA to bind SspB leading eventually to the functional activation of OptoKANK. **(B)** Representative images of OptoKANK dimerization in HT1080 cells upon photoactivation. Before illumination, KANK-ΔKN-mEmerald (ΔKN) is diffused within the cell. After 150 sec of blue light illumination, ΔKN is recruited to KN. The graphs show the KN and ΔKN mean intensities over the line (dotter white line) in the representative images, before and after photoactivation of KANK. **(C)** Representative images of Vinculin-mIFP-transfected HT1080 cells carrying the full OptoKANK constructs (KN + ΔKN), or either the KN construct or the ΔKN construct, before and after 8 min of blue light illumination on the focal adhesion (blue dotted circle). Graph shows the average of mean vinculin normalized as a fraction of initial intensity of illuminated focal adhesion over the time in these three conditions. (Data are presented as mean ± s.e.m; n = 19 minimum). **(D)** Representative images of Vinculin-mIFP-transfected HT1080 cells carrying the full OptoKANK constructs (KN+ ΔKN) before and after 8’ of blue light illumination on the focal adhesion (blue dotted cercle) for cells in control (Ctrl DMSO), treated with 1 µM of nocodazole, and after the nocodazole washout. Graph shows the vinculin intensities normalized as a fraction of initial intensity of illuminated focal adhesions over the time in these three conditions. (Data are presented as mean ± s.e.m; n = 8 from two independent experiments). **(E)** Time-course of the focal adhesion disassembly after nodazole washout upon OptoKANK activation (blue dotted circle).

Assessment of the vinculin fluorescence intensity revealed that blue illumination of focal adhesion in cells containing complete OptoKANK (KN + ΔKN) induced decrease of vinculin density accompanied by centripetal sliding of the focal adhesion (**figure 1C** **+ supp. movie 1)**. This resulted in almost total disappearance of the adhesion within about 10 minutes. The focal adhesions in cells expressing incomplete OptoKANK (either only KN or ΔKN) were not affected by the illumination (**figure 1C**). To check whether this effect was due to targeting of microtubules to focal adhesions we compared the effect of focal adhesion illumination in non-treated cells expressing OptoKANK with that in cells treated with nocodazole for 2 hours and cells treated with nocodazole and then washed out for 2 hours. We found that disruption of microtubules with nocodazole abolished the effect of illumination on the focal adhesion integrity while washing the drug out restored the disruptive effect of illumination (**figure 1D** **and 1E + supp. movie 2 and movie 3)**. Thus, the optogenetics driven formation of KANK1-mediated link between microtubules and focal adhesions resulted in sliding and disassembly of focal adhesion in a microtubule dependent fashion.

### Targeting of microtubules to the focal adhesion area upon local OptoKANK activation

We further assessed how the number of microtubule tips in the focal adhesion area changes upon local photoactivation of OptoKANK. In the first series of experiments, the microtubule tips were labeled with EB3-mIFP co-transfected with the OptoKANK constructs. The focal adhesions were identified by labelled KN domain (mEmerald-KANK1-LOV2ssrA), which as mentioned above, always co-localized with vinculin-labelled focal adhesion (**figure 1B**). Manual counting of EB3 “comets” overlapping with focal adhesion area revealed approximately a 25% increase in the number of microtubule tips already at 30 seconds following the onset of illumination (**figure 2A**). There were no changes in the average number of microtubule tips at non-illuminated focal adhesions in the same cell during this time interval (**figure 2A**).

**Figure 2:**
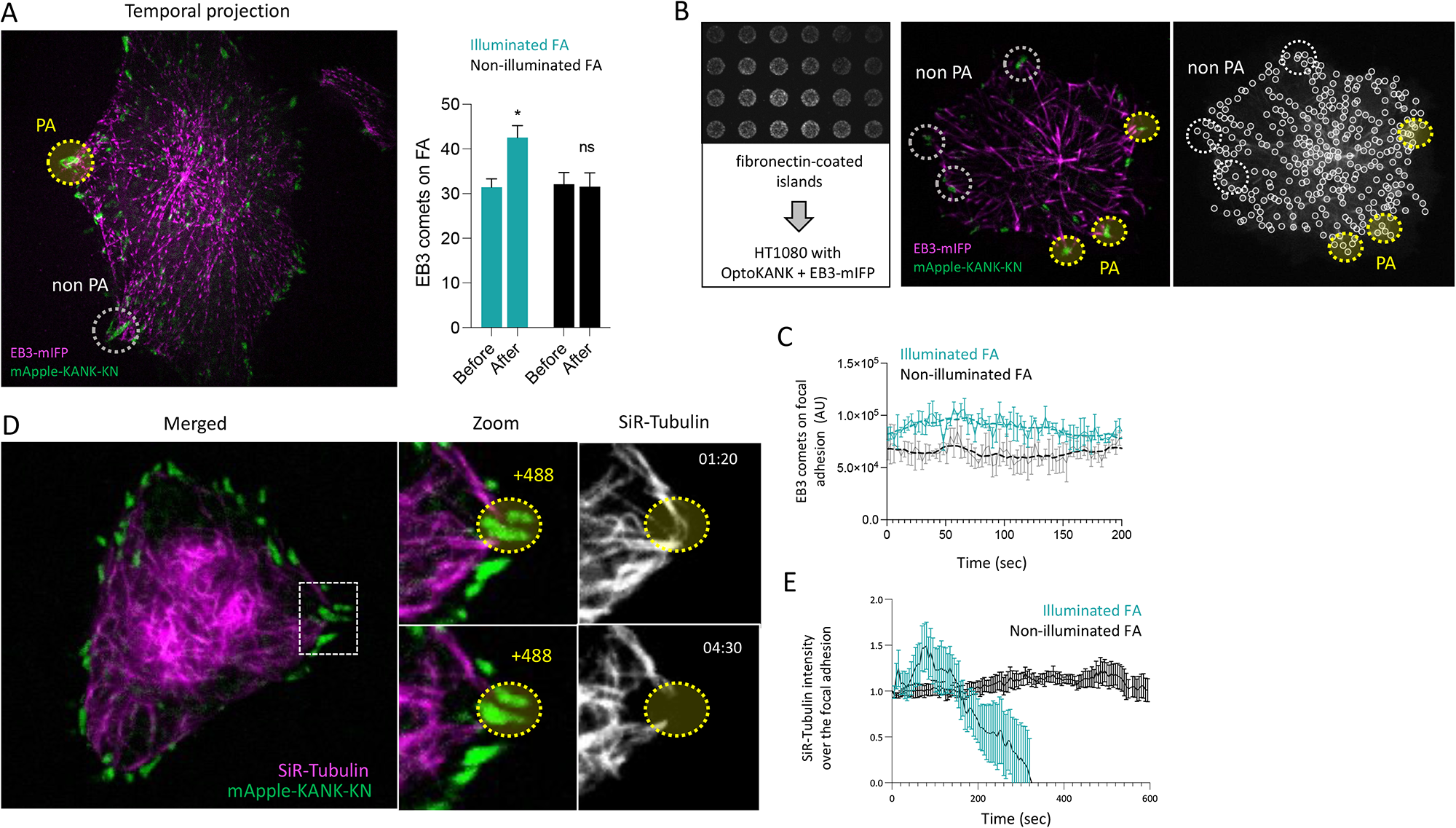
Augmented targeting of microtubules to the focal adhesion area upon local OptoKANK activation. **(A)** Temporal projection of EB3-mIFP in OptoKANK-transfected HT1080 cells. (1) and (2) are zooms of photoactivated (PA – yellow dotted circle) and non-photoactivated (non PA – grey dotted circle) focal adhesion respectively. Focal adhesions are visualized with KN construct. Graphs show the average number of EB3 comets reaching the illuminated and non-illuminated focal adhesions before and after 30 sec of blue light illumination. (Data are presented as mean ± s.e.m; n = 9). **(B)** Representative image of fibronectin-coated islands on which the HT1080 cells transfected with OptoKANK and EB3-mIFP were plated. EB3-mIFP marker was used to assess the number of EB3 comets that reach photoactivated (PA – yellow doted circles) and non-photoactivated (non PA – white dotted circles) focal adhesions which were visualized using the KN construct. On the right panel, representative image processed with the U-Track 2 script (see material and methods) that was used to quantify the number of EB3 comets reaching the different regions of interest. **(C)** Graph shows the integrated EB3 comet fluorescence after processing with U-Track2 for photoactivated and non-photoactivated focal adhesions over the time. (Data are presented as the mean of integrated EB3 fluorescence from 3 focal adhesions under each condition). **(D)** Representative image of OptoKANK-transfected HT1080 cell after 1 min 20 sec and 4 min 30 sec of blue light illumination of the focal adhesion (dotted yellow line). SiR-Tubulin (magenta) at 250 nM added 3 hours prior the experiments was used to visualize microtubule. **(E)** Graph shows the mean SiR-Tubulin intensity at the photoactivated and non-photoactivated focal adhesions over the time (Data are presented as mean ± s.e.m; n = 18 from 2 independent experiments).

To obtain further quantitative data, we plated cells on a micropatterned substrate with circular fibronectin-coated islands of 4.5 µm in diameter. On this substrate cells could only form focal adhesions within these adhesive islands (usually one to three focal adhesion per island – **supp. figure 1A**). We further measured the dynamics of fluorescence intensity of EB3-mIFP over time in both illuminated and non-illuminated islands. We illuminated three islands per cell leaving others non-illuminated. The analysis of the time course of EB3 fluorescence revealed an apparent variability in both illuminated and non-illuminated areas (**Figure 2B and 2C** + **supp. figure 1B + supp. movie 4**). However, in all cases the fluorescence intensity of EB3 per island was higher in the illuminated islands with photoactivated OptoKANK (**Figure 2B and 2C + supp figure 1B**) consistent with measurements shown in **figure 2A**.

The EB1 and EB3 proteins recognize the growing plus ends of microtubules that contain GTP caps but not the shrinking ends^27^. Therefore, to fully quantify the microtubules targeting to focal adhesions, we used SiR-Tubulin labeling^28^. These experiments revealed a stereotypic dynamic of the intensity of SiR-Tubulin fluorescence overlapping with focal adhesions upon OptoKANK activation by illumination. The SiR-Tubulin fluorescence increased in the first 60 seconds following onset of illumination and then decreased below the level typical for non-illuminated focal adhesions (**figure 2D** **and 2E**). These results suggest that activation of OptoKANK first promoted targeting of microtubules to focal adhesions and then synchronous microtubule withdrawal.

The microtubules recruited to the focal adhesions by OptoKANK illumination preserved their transport functions. As it has been demonstrated in some cellular systems, microtubules deliver endocytic vesicles to the plasma membrane-associated secretory sites in a KANK dependent fashion^21, 29^. Normally, accumulation of the new membrane at these sites is balanced by the local endocytosis. We expected that in the presence of endocytosis inhibitor, dynasore^30^, the accumulation of the functional microtubule tips by OptoKANK illumination will augment the local delivery of the new membrane. Indeed, we have found that illumination of focal adhesions in cells containing OptoKANK construct treated with dynasore resulted in formation of membrane blebs at the illuminated sites **(supp. figure 1C + supp. movie 5).**

### Dynamics of myosin-II filaments and traction forces upon local OptoKANK activation

Observations of myosin-II filaments labeled with myosin light chain fused with near-infrared fluorescent protein (MLC-iRFP) revealed that activation of OptoKANK by local illumination of focal adhesions resulted in rapid accumulation of myosin filaments in the proximity of focal adhesions (**figure 3A** **+ supp. movie 6**). Myosin-II filaments fluorescence signal appeared in a centripetal direction from the focal adhesions with a maximum at approximately 4 µm distance from the focal adhesion proximal end (**figure 3B**). Time course analysis of the burst of myosin-II filaments accumulation revealed that the myosin filament density approaches the maximum in about 100 seconds following the onset of illumination (**figure 3C** **and 3D)**. The accumulation of F-actin visualized by SiR-Actin^27^ was observed at the same time and location **(supp. figure 2A + supp. movie 7)**. Comparison of the dynamics of myosin filaments accumulation with those of microtubule and vinculin density in focal adhesion area showed that the appearance of myosin filaments at the proximal end of focal adhesion coincided with shrinking of microtubules that were associated with focal adhesions at the onset of illumination (**figure 3D**). In turn, the moment when the density of myosin filaments approached the maximum preceded the process of focal adhesion disassembly (**figure 3D**).

**Figure 3:**
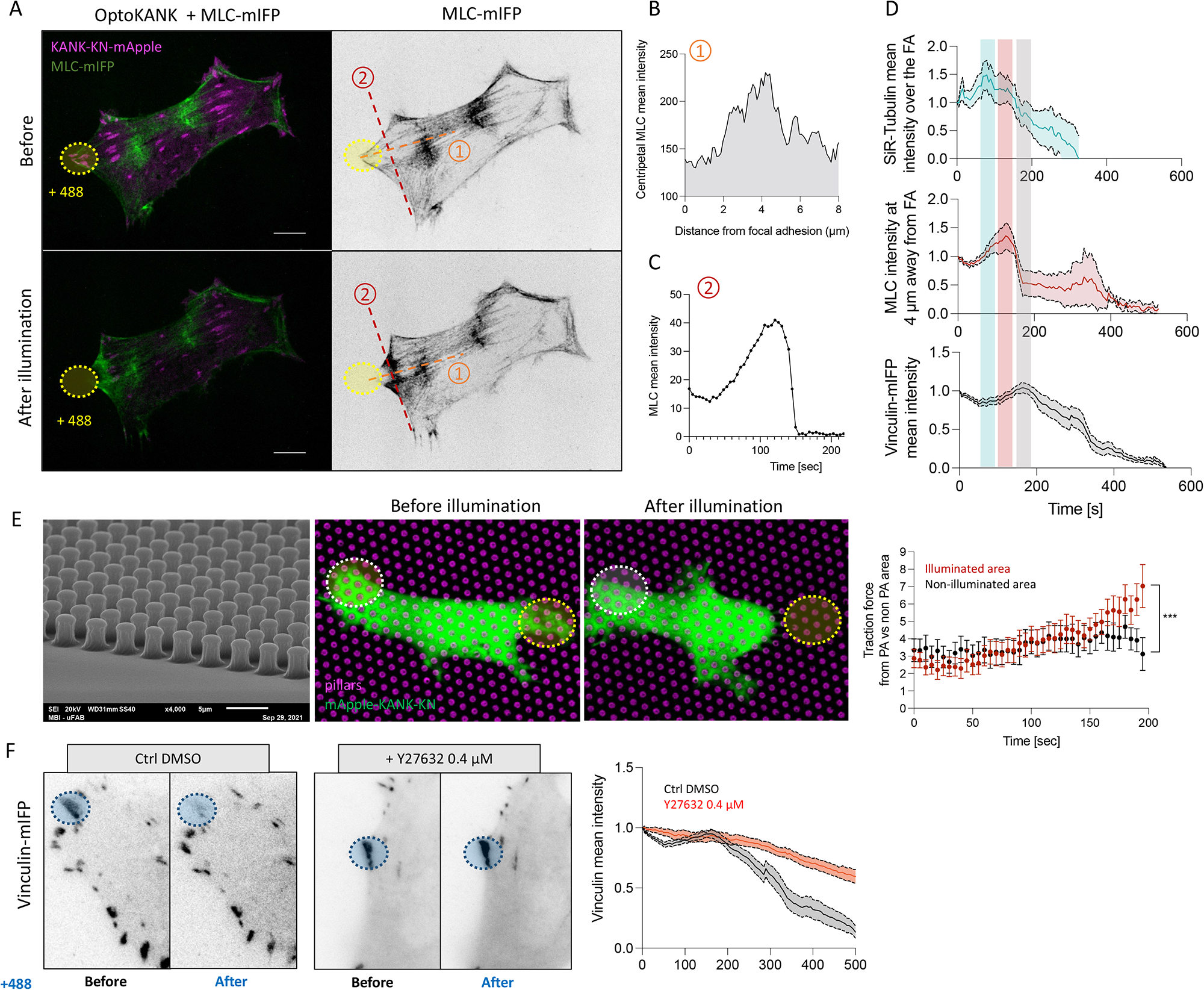
Dynamics of myosin-II filaments and traction forces upon local OptoKANK activation. **(A)** Representative image of OptoKANK-transfected HT1080 cell before and after 120 sec of blue light illumination on the focal adhesion (yellow dotted line). MLC-mIFP was used to assess the myosin-II dynamics upon OptoKANK activation. The line scan (1) was used to measure the MLC-mIFP intensity in the vicinity (4 µm away in centripetal direction) of the proximal end of the photoactivated focal adhesion shown in **(B)**. The line scan (2) was used to measure the MLC-mIFP mean intensity in the proximity of the photoactivated focal adhesion over the time **(C)**. **(D)** Time courses of the microtubule intensity overlapping with focal adhesions (SiR-Tubulin – blue rectangle), MLC intensity at 4 µm from the proximal end of the illuminated focal adhesion (MLC-mIFP mean intensity – red rectangle) and vinculin mean intensity in focal adhesion (Vinculin-mIFP mean intensity – grey rectangle). Note that increase of the microtubule intensity upon illumination succeeded by its drop, which coincide with the increase of MLC intensity in proximity of the focal adhesion which preceded the drop in vinculin intensity. (Data are presented as mean ± s.e.m; Ctrl, n = 8; MLC intensity, n = 6; SiR-Tubulin, n = 18. Data are from at least two independent experiments). **(E)** Representative images of OptoKANK-transfected HT1080 cell before and after blue light illumination on the yellow area (yellow dotted circle). The cells were plated on micropillars (magenta) for traction force assessment over the time. Graph shows the average traction force at the focal adhesion surrounding (based on measurement of deflection of about 3-4 pillars per focal adhesion) upon illumination-induced OptoKANK activation compared to the traction forces in non-illuminated area. (Data are presented as mean ± s.e.m; n = 19 pillars from 6 different cells). **(F)** Representative images of Vinculin-mIFP-transfected HT1080 cells carrying the OptoKANK constructs (KN + ΔKN) before and after 8 min of blue light illumination of the focal adhesion (yellow dotted line) for control cells (Ctrl DMSO), and cells treated with 0.4 µM of Y27632. Graph shows the normalized mean vinculin intensity of illuminated focal adhesions over the time under these two conditions. (Data are presented as mean ± s.e.m; n = 18 cells minimum from two independent experiments).

To check if accumulation of myosin filaments affects the traction forces applied through the focal adhesions, we plated cells on the array of elastic micropillars^31^ and assessed the time-course of micropillar deflections in both illuminated and non-illuminated areas (**figure 3E**). Measurements of traction force at the focal adhesion area (approximately 4 pillars per focal adhesion) over the time revealed a significant increase of traction force upon illumination-induced OptoKANK activation compared to the non-illuminated area (**figure 3E**).

Accumulation of myosin filaments in proximity to focal adhesions and development of traction forces appeared to be critically important for the disassembly of focal adhesions upon forced targeting of microtubules by OptoKANK activation. Treatment with Y27632 abolished the traction force increase and prevented the sliding and disassembly of focal adhesions (**Figure 3F**). Of note, the treatment with Y27632 in these experiments was relatively mild and short, lasting only for the period of illumination. Y27632 was used only to prevent the increase of traction forces developed due to myosin II filaments accumulation upon microtubule targeting to focal adhesions. More pronounced treatment with these drugs which resulted in complete annulation of traction forces led to disassembly of focal adhesions as was well established in numerous previous studies^9, 32^.

Thus, targeting of microtubules to focal adhesions by OptoKANK activation results in transient accumulation of myosin-II filaments near the proximal end of the focal adhesions and bursts of traction force, which are required for focal adhesion disassembly.

### Rho activation by GEF-H1 is required for focal adhesion disassembly upon microtubule targeting

Disruption of microtubules or their disconnection from integrin adhesions induces formation of myosin filaments due to release and activation of GEF-H1 and subsequent activation of Rho and ROCK^15, 16^. Thus, we decided to check whether transient activation of myosin filaments formation in the proximity of focal adhesions and subsequent focal adhesion sliding and disassembly upon OptoKANK activation also depends on GEF-H1 and Rho. We found that in GEF-H1 depleted cells activation of OptoKANK by the illumination of focal adhesions did not result in increase of traction force applied to the focal adhesions (**figure 4A**). Consistently the OptoKANK activation in GEF-H1 depleted-cells did not trigger the sliding and disassembly of the focal adhesions (**figure 4B** **+ supp. movie 8**).

**Figure 4:**
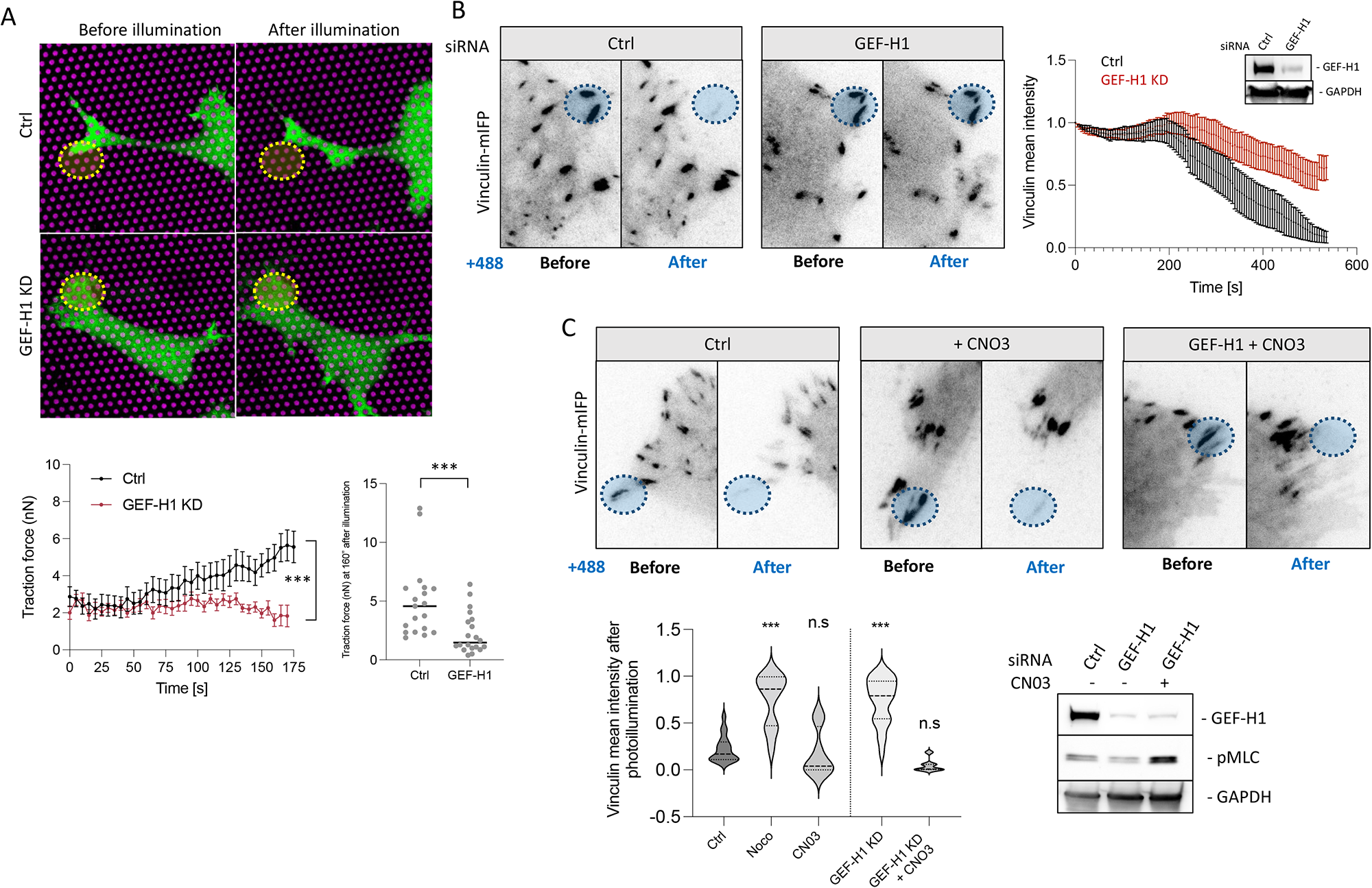
Rho activation by GEF-H1 is required for focal adhesion disassembly upon microtubule targeting. **(A)** Representative images of OptoKANK-transfected HT1080 cell before and after blue light illumination on yellow area (yellow dotted circle) and plated on micropillars (magenta) for traction force assessment over the time in control (Ctrl) or GEF-H1 knockdown (GEF-H1 KD) conditions. Graph bellow shows the traction force of pillars in the region of interest over the time for control (CTRL) or GEF-H1 depleted cells (GEF-H1 KD). Traction force after 160 sec of blue light illumination is shown on the right panel (Data are presented as mean ± s.e.m; Ctrl, n = 19 from 6 different cells; GEF-H1 KD, n = 20 from 7 different cells). **(B)** Representative images of Vinculin-mIFP-transfected HT1080 cells carrying the OptoKANK constructs before and after 8’ of blue light illumination on the encircled focal adhesion (yellow dotted line) for cells in control (Ctrl), or depleted for GEF-H1 (GEF-H1 KD). Graph shows the normalized mean vinculin intensity of illuminated focal adhesions over the time in these two conditions (Data are presented as mean ± s.e.m; n = 20 minimum from three independent experiments). Immunoblots of GEF-H1 and GAPDH in ctrl and GEF-H1 depleted cells are shown in the black box. **(C)** Representative images of Vinculin-mIFP-transfected HT1080 cells carrying the OptoKANK constructs before and after blue light illumination on the encircled focal adhesion (yellow dotted line) for control cells (Ctrl), cells treated with CNO3 and for GEF-H1 depleted cells treated with CNO3 (GEF-H1 KD + CN03). Graph shows the normalized mean vinculin intensity after the illumination for control cells (CTRL), cells treated with 1 µM nocodazole, cells treated with CNO3, and GEF-H1 depleted cells treated with CN03 (Data are presented as mean ± s.e.m; Ctrl, n = 18; Noco; n = 12; CNO3, n = 8; GEF-H1 KD, n=24; GEF-H1 KD + CNO3, n=7. Data are from two intendent experiments). Immunoblots of GEF-H1, pMLC and GAPDH in these three conditions are in the black box.

Further, we rescued the effect of GEF-H1 knockdown by activation of Rho. Treatment of OptoKANK expressing cells with Rho activator CNO3 increased the focal adhesion sizes but did not prevent the sliding and disassembly of focal adhesions upon OptoKANK activation by illumination (**figure 4C**). The treatment of OptoKANK expressing GEF-H1 knockdown cells with CNO3 during the focal adhesion illumination rescued the effect of GEF-H1 knockdown and resulted in sliding and disassembly of focal adhesions (**figure 4C** **+ supp. movie 9**). Thus, the burst of traction force and consequent disassembly of focal adhesions upon microtubule targeting depends on Rho activation by GEF-H1.

### OptoKANK mediated disassembly of focal adhesions requires activity of FAK, PAK, Kinesin-1, αTAT1 and APC

Targeting of microtubules to focal adhesions by local activation of KANK provides a convenient experimental system to elucidate the function of different proteins in the microtubule-driven focal adhesion disassembly. As shown above, GEF-H1 and ROCK are important players in this process. Using siRNA mediated knockdowns or/and pharmacological inhibitors, we screened a few candidates and established the involvement of several microtubule- and focal adhesion-associated proteins in the focal adhesion disassembly process induced by activation of KANK1-mediated link. The majority of the candidates included in our screen were based on previous publications suggesting the involvement of these proteins in the microtubule driven regulation of focal adhesion turnover^8, 22, 25, 33–41^. The graph summarizing the results of these experiments is shown in **figure 5A** and **supplementary figure 3A**. The readout in these experiments was the alteration of vinculin fluorescence intensity of the focal adhesions in cells expressing OptoKANK constructs upon illumination of this adhesion for 10 minutes. Among the 18 proteins screened, 7 were needed for the disassembly of focal adhesions induced by OptoKANK activation. Besides GEF-H1 and ROCK (**figure 3F** **and 4B**), the 5 proteins identified are the focal adhesion kinase FAK, the PAK family kinase members sensitive to FRAX1036 inhibitor^42^, kinesin-1 as suggested by experiments with kinesore inhibitor^43^, tubulin acetylase αTAT1 and microtubule associated actin regulator adenomatous polyposis coli (APC) protein (**Figure 5A**). The results implicating FAK, APC and Kinesin-1 in microtubule-driven focal adhesion disassembly are consistent with previous publications^25, 33, 44^.

**Figure 5:**
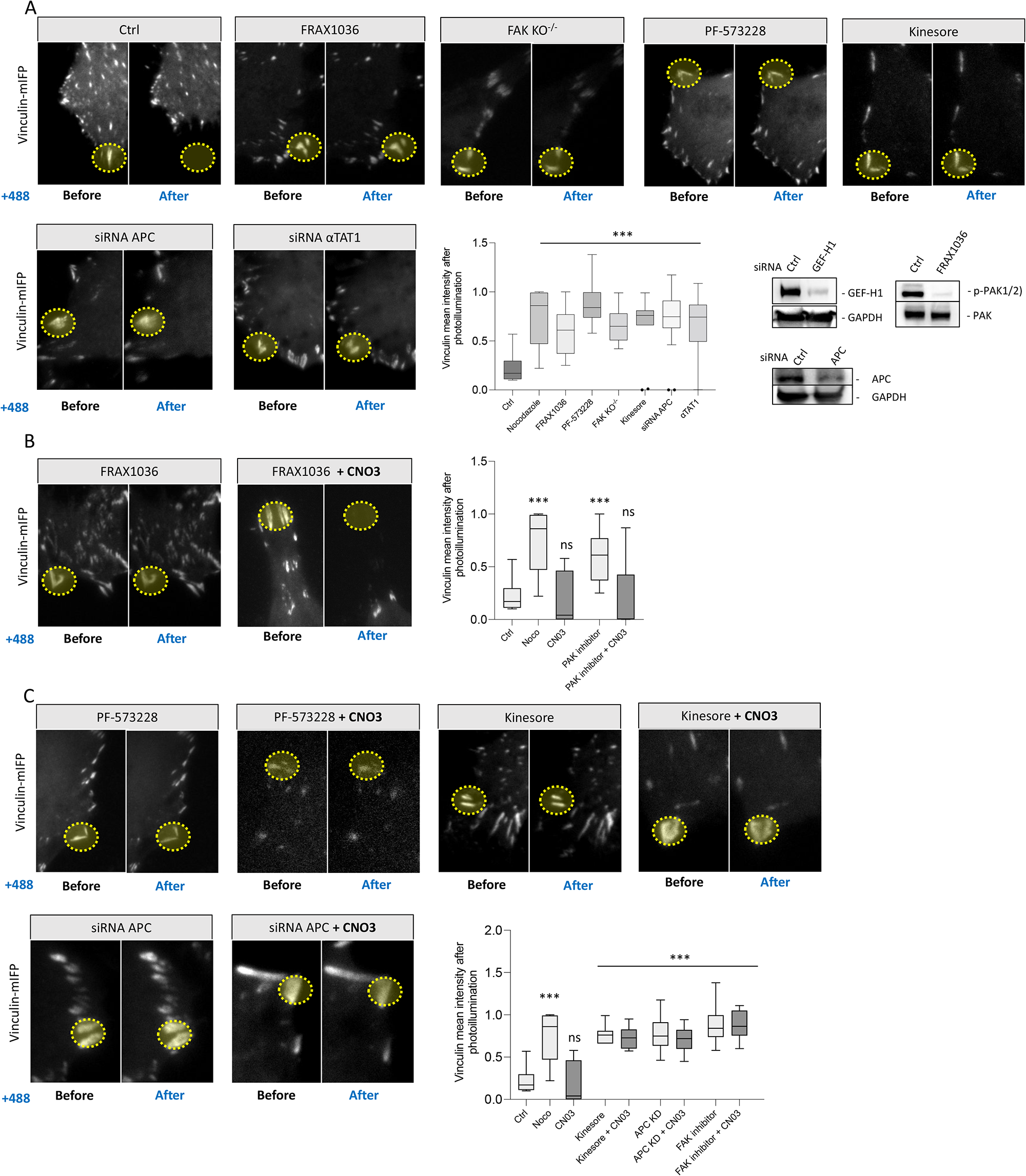
OptoKANK mediated disassembly of focal adhesions requires activity of FAK, PAK, Kinesin-1, αTAT1 and APC. **(A)** Representative images of Vinculin-mIFP-transfected HT1080 cells carrying the OptoKANK constructs before and after blue light illumination of the focal adhesion (yellow dotted line) for control cells (Ctrl), cells treated with FRAX1036, PF-57328, Kinesore, as well as in APC-depleted, αTAT1-depleted cells, and in FAK^-/-^ knockout MEF cells. Graph shows the normalized mean vinculin intensity after the illumination of cells treated as indicated (Data are presented as mean ± s.e.m; Ctrl, n =18; Nocodazole, n = 12; FRAX1036, n = 15; PF-573228, n = 10; FAK KO^-/-^, n = 8; Kinesore, n = 11; siRNA APC, n = 18; αTAT1, n = 10). Immunoblots of GEF-H1, p-PAK1/2, APC and GAPDH are shown in the black box. **(B)** Representative images of Vinculin-mIFP-transfected HT1080 cells carrying the OptoKANK constructs before and after blue light illumination on the focal adhesion (yellow dotted line) for cells treated with FRAX1036, and cells treated with FRAX1036 simultaneously with the contractility activator CNO3. Graph shows the normalized mean vinculin intensity after the illumination for cells under these conditions (Data are presented as mean ± s.e.m; Ctrl, n = 18; Noco, n = 12; PAK inhibitor, n = 15, PAK inhibitor + CN03, n = 10). **(C)** Representative images of Vinculin-mIFP-transfected HT1080 cells carrying the OptoKANK constructs before and after blue light illumination of the focal adhesion (yellow dotted line) for cells treated with FAK inhibitor PF-57328, Kinesine-1 modulating drug Kinesore, and APC-depleted cells and in such cells treated with Rho activator CNO3. Graph shows the normalized mean vinculin intensity after the illumination for cells under treatments mentioned above (Data are presented as mean ± s.e.m; Ctrl, n =18; Nocodazole, CNO3, n = 8; Kinesore, n = 10; Kinesore + CN03, =n = 16; APC KD, n = 18; APC KD + CNO3, n = 14, FAK inhibitor; n = 10; FAK inhibitor + CNO3, n = 10).

The suppressive effect of GEF-H1 knockdown on focal adhesion disassembly by OptoKANK activation can be cancelled by activation of Rho (**figure 4C**), which suggests that the function of GEF-H1 in this process is indeed RhoA activation. To elucidate which other proteins among the 7 candidates identified by our screen are involved in Rho/myosin IIA activation after microtubule targeting, we checked whether activation of Rho by CNO3 can cancel the inhibitory effect of knockdowns/pharmacological inhibition of these proteins. We found that besides GEF-H1, the effect of PAK kinase inhibition can be abolished by CNO3 treatment (**figure 5B**) suggesting that PAK function in microtubule driven focal adhesion disassembly is also related to Rho/myosin II activation in agreement with previous publication on PAK function^45^. Effects of knockdown or/and pharmacological inhibition of other candidates (FAK, Kinesin-1, APC) were not rescued by CNO3 treatment (**figure 5C**) suggesting that these proteins are involved in different stages of the process of microtubule-driven focal adhesion disassembly.

### Working hypothesis and a physical model of the process of OptoKANK activation driven focal adhesion disassembly

Our results suggest that local activation of actomyosin contractility in the proximity of focal adhesions is a necessary step in the OptoKANK activation-driven disassembly of a focal adhesion. Indeed, we have shown that this process depends on GEF-H1 and ROCK and detected accumulation of actomyosin and increase of traction force in the proximity of a focal adhesion preceding its disassembly. In the absence of microtubule targeting the augmentation of force applied to the focal adhesion results in their growth rather than disassembly^9^, therefore we assume that targeting of microtubules to the focal adhesion weakens the adhesion, enabling the focal adhesion to slide upon activation of myosin driven traction force. During this sliding, the force experienced by the focal adhesion drops^46^, triggering its disassembly. Thus, based on our results, the process of focal adhesion disassembly after OptoKANK activation consists of the following steps (**figure 6A-D**). i) Accumulation of microtubule tips at focal adhesions triggering weakening of the adhesions but not yet their disassembly (**figure 6B**); ii) the detachment and withdrawal of microtubule tips accompanied by release and activation of GEF-H1 which, with the assistance of PAK, results in accumulation of actin and myosin II filaments in the vicinity of the proximal end of the focal adhesion and activation of traction force (**figure 6C**); iii) detachment and sliding of the focal adhesion as a result of burst of traction force leading to its disassembly (**figure 6D**).

**Figure 6:**
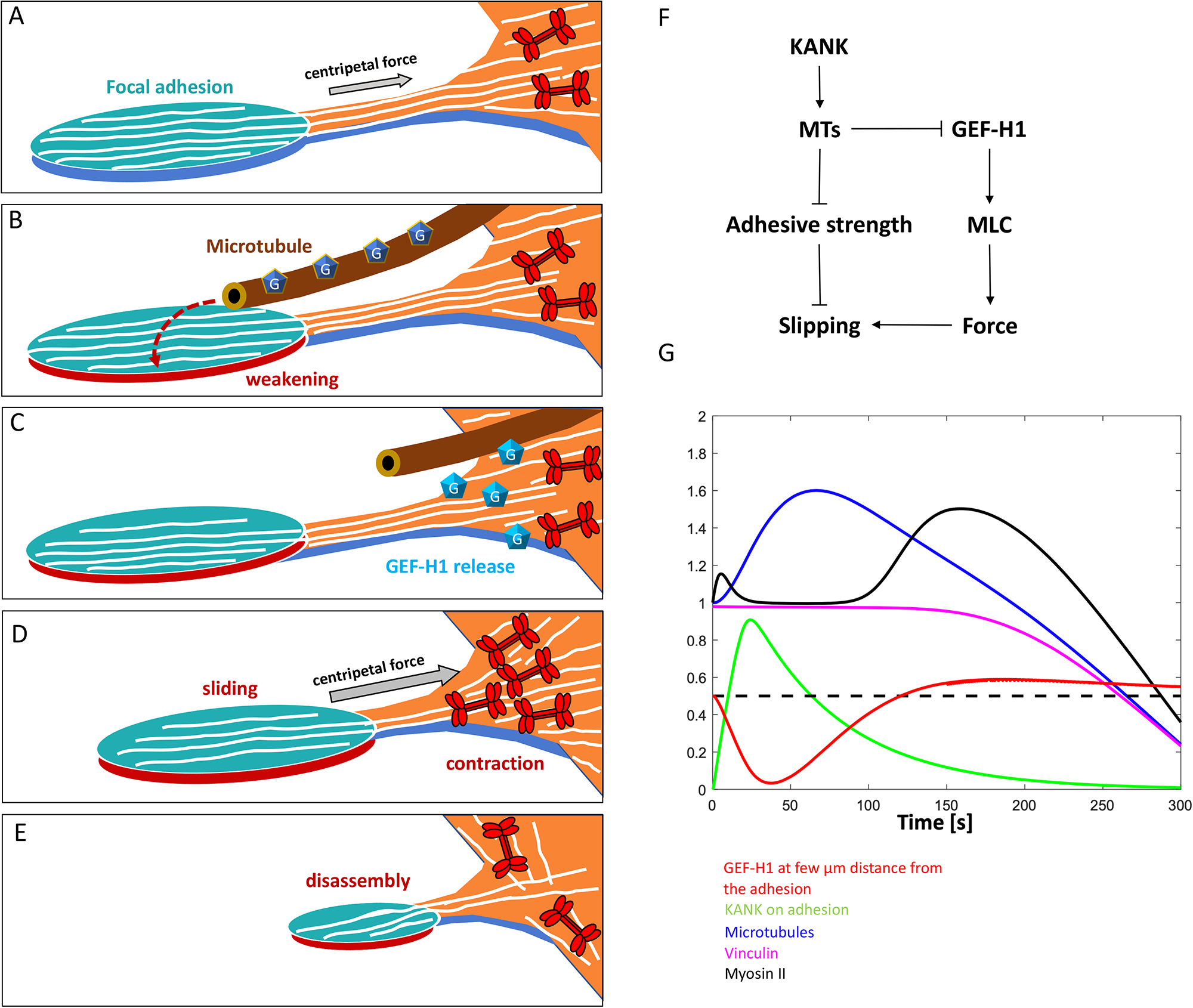
Working hypothesis and a physical model of the process of OptoKANK activation driven focal adhesion disassembly. **(A-D)** Cartoons depicting the proposed model of sequential events leading to the microtubule-driven focal adhesion disassembly upon OptoKANK activation. **(A)** Focal adhesion and associated stress fiber embedded in contractile actomyosin network are depicted. Actin filaments are schematically presented by white lines and myosin II filaments by red symbols (not in scale). The focal adhesion is stabilized by centripetal force generated by the actomyosin network. **(B)** Microtubule attachment to focal adhesion weakens it but the centripetal force is not yet sufficient to promote the sliding. Microtubule is associated with non-active GEF-H1 molecules (dark blue pentagons with “G” inside). **(C)** Microtubule detachment from the focal adhesion triggers the release of GEF-H1 and its activation in the vicinity of the proximal end of the focal adhesion (represented by changes in the color of the symbols to light blue). **(D)** Active GEF-H1 triggers the Rho/ROCK pathway resulting in formation of new myosin and actin filaments and local actomyosin contraction. The strong centripetal force developed as a result of such contraction leads to sliding of weakened focal adhesion. **(E)** Sliding of the focal adhesion leads to it disassembly. **(F)** Diagram showing the feedbacks included into the physical model. **(G)** Model-predicted time series for dynamics of microtubules, vinculin, KANK, GEF-H1 and myosin-II upon optogenetic induction of microtubules contact with the focal adhesion (see text). Model parameters used in the simulations are described in the **Supplemental Methods**.

To analyze whether this working hypothesis can explain the process of microtubule-driven focal adhesion disassembly in accordance with experimental results, we propose a simple physical model of this process. In the model, we simulate on the computer dynamics of a) number of microtubules in contact with a focal adhesion, b) number of KANK molecules on the focal adhesion, c) myosin II filament density proximal to the focal adhesion, d) adhesion molecule density gripping the substrate, and e) density of active GEF-H1 molecules in the cytoplasm proximal to the focal adhesion. The model is based on the following assumptions: 1) growing microtubules arrive to the focal adhesion at a constant rate and pause there before shortening and leaving the adhesion. Based on the comparison of the microtubule dynamics at the cell periphery in control and KANK1/2 depleted cells^18^, it is reasonable to suggest that the average pause is an increasing function of the number of KANK molecules on the focal adhesion. Since the effects of microtubules on focal adhesion depends on kinesin-1 (ref ^33^ and this study), microtubules may deliver to focal adhesions some factors which facilitate dissociation of integrins and talins, and because KANK binds to talin, of KANK molecules themselves. Indeed, the fluorescence intensity of talin-1, integrin beta3, and KANK-KN domain started to decrease after 30-60 seconds of illumination, much earlier than vinculin **(supplemental figure 4A)**. 2) Based on the known affinity of GEF-H1 to microtubules^15^, we assume that any time a microtubule arrives at the focal adhesion, a certain number of GEF-H1 molecules are locally absorbed from the cytoplasm; when this microtubule leaves the focal adhesion, after a pause, GEF-H1 molecules are released back into the cytoplasm, where they diffuse and undergo activation^47^. Rates of myosin activation increases when active GEF-H1 concentration in the cytoplasm is above a threshold. To not overwhelm the model, we do not explicitly include steps of activation of the released GEF-H1 molecules or ROCK-dependent pathway of myosin activation. In agreement with our observations, we assume in our model that activation of GEF-H1 and myosin-II occurs in the centripetal direction from focal adhesion, in the region where the microtubule tips retreated from the focal adhesion are located. 3) We assume that the pulling force applied to the focal adhesion is directly proportional to the myosin density in the model and that the focal adhesion is gripping the substrate if the ratio of the pulling force to adhesive strength is below a threshold, and slipping if this ratio is above the threshold. The slipping of focal adhesion triggers its rapid disassembly^46^ (as manifested by the drop in vinculin concentration). Finally, 4) we assume that the adhesive strength is proportional to the KANK density on the focal adhesion (not that KANK directly contributes to the adhesive strength, but rather that its adhesive molecular partner talin does, and that talin amount is proportional to that of KANK). The feedbacks included into the model are shown in **figure 6F**. Model details, parameters and variations are discussed in the supplemental text.

The predicted time series of the key molecular densities upon local activation of KANK shown in **figure 6F** compare well with the experimental data. The model predicts that up to ∼ 60 sec after the illumination, microtubules arriving to the focal adhesion locally sequester GEF-H1 molecules, sharply depleting GEF-H1 density near the focal adhesion, but after that the decreasing microtubule number leads to release of the accumulated GEF-H1 molecules. If the GEF-H1 dynamics was strictly local, then GEF-H1 molecules would be first captured by the microtubules, then released in the same place, with no net gain. Crucially, due to diffusion, and also because the sink for GEF-H1 on the increasing number of microtubules occurs earlier than the source of GEF-H1 from the decreasing number of microtubules, GEF-H1 molecules initially diffuse closer to the focal adhesion, down the gradient created by the sink. After that, when GEF-H1 is released by disassembling microtubules, the net GEF-H1 concentration increases after a short delay in the vicinity of the proximal end of the focal adhesion. Effectively, the initial microtubule number increase, counter-intuitively, helps by ‘soaking’ GEF-H1 into the focal adhesion vicinity, then releasing increased amounts of GEF-H1. Because this diffusion-reaction process introduces delays, GEF-H1 concentration becomes greater than the baseline ∼ 120 sec after the local activation of KANK. This leads to a significant additional activation of myosin and traction force increase by ∼ 150 sec after the local activation of KANK. By that time, the adhesion is significantly weakened, the force to adhesive strength ratio exceeds the gripping-to-slipping threshold, and the biphasic process comprising of weakening of adhesion to the substrate and local GEF-H1/RhoA/ROCK dependent activation of contractility leads to slippage (**figure 6F**). The model also correctly accounts for the results of several perturbation experiments (**Supplemental Figure 5A-D**). The model predicts that: 1) Local KANK activation in GEF-H1 KD cell does not result in the traction force increase, hence no adhesion sliding (**Supplemental Figure 5A).** 2) In cells with Rho activated by CNO3, either with or without functional GEF-H1, there is an elevated pulling on the focal adhesion before the activation of KANK, but the force to adhesion strength ratio is still below the threshold that causes sliding. After the activation of KANK, microtubules’ arrival brings adhesion weakening, which, together with the elevated pulling, triggers the sliding (**Supplemental Figures 5B and 5C).** 3) When the ROCK activity is weakened by Y27632, despite the Rho activity increase due to the GEF-H1-dependent activation, myosin-II-dependent pulling force is too weak to slide the adhesion even against the weakened adhesion (**Supplemental Figure 5D)**.

## Discussion

In this study, we used a novel optogenetic method to decipher the process of microtubule-driven focal adhesion disassembly. We have shown that transient attachment of microtubule tips to a focal adhesion followed by their detachment results in a local burst of actomyosin contractility, which is a critically important step in the process that triggers focal adhesion sliding and consequently its disassembly.

The optogenetic construct connecting microtubule tips with focal adhesions consisted of two halves of KANK1 protein, talin binding domain (KN) and the rest of the molecule (ΔKN) that can be linked by illumination. Since these constructs were overexpressed as compared to endogenous KANK, they likely displaced the endogenous KANK from the focal adhesions and thereby uncoupled the KANK-mediated link between focal adhesions and microtubules. This scenario is supported by the fact that the adhesion area in cells overexpressing OptoKANK constructs was significantly larger than that in mock transfected cells, similarly to the situation in cells with disrupted microtubules or following KANK1 knockdown^16^. We have shown that local illumination restoring the integrity of the KANK molecule resulted in transient increase of microtubule tips overlapping with focal adhesions and augmentation of the delivery of membrane to the focal adhesion area. Further, optogenetic targeting of microtubules leads to the disassembly of focal adhesions similarly to that observed in cells recovering following washout of microtubule disrupting drugs^25^. Thus, our optogenetic approach permitted us to investigate in detail the kinetics of microtubule-driven focal adhesion disassembly and identify the major molecular players participating in this process.

The key observation was that the initial increase of the number of microtubules associated with focal adhesion was followed by withdrawal of microtubules from the focal adhesion zone which in turn resulted in sliding and gradual disassembly of the focal adhesion. Our previous studies suggested that detachment of microtubules from integrin adhesions triggers the release of GEF-H1 from microtubules and subsequent formation of myosin-II filaments^16^. In agreement with this idea, we have shown here that (i) GEF-H1 is critically important for triggering microtubule-driven focal adhesion sliding and disassembly; (ii) local assembly of myosin-II and actin filaments is observed near the proximal end of focal adhesion after microtubule retraction; (iii) inhibition of acto-myosin contractility by Y27632 prevents the sliding and disassembly of illuminated focal adhesions; (iv) activation of Rho by CNO3 restores the disassembly of focal adhesion upon illumination in GEF-H1 knockdown cells. Altogether, these data suggest that local retraction of microtubules after their accumulation above a focal adhesion upon OptoKANK activation results in release and activation of GEF-H1, which in turn triggers activation of Rho. This activation leads to local formation of an actomyosin contractile network in proximity to the focal adhesion, and development of traction forces pulling the adhesion in a centripetal direction. Furthermore, we detected the increase of traction force at the focal adhesion area that occurs after microtubule retraction and showed that it preceded the sliding and disassembly of focal adhesion.

In our model, the focal adhesion disassembly occurs as a result of focal adhesion sliding. Indeed, focal adhesions are mechanosensitive in a sense that they undergo rapid disassembly in the absence of stretching force generated by actomyosin contractility^6–11^. Detached and sliding focal adhesions should not experience any significant stretching force and therefore should undergo rapid disassembly similar to that induced by inhibition of myosin contractility^46^. Based on the data summarized above, we assume that the main factor inducing focal adhesion sliding is the local burst of actomyosin contractility in the proximity of the focal adhesion which we observed in our experiments. However, the increase of contractility alone does not seem to be sufficient for detachment and sliding of focal adhesion. As mentioned above, increase of contractility in cells attached to rigid substrate covered with integrin ligands results in an increase of focal adhesion size, due to the mechanosensory nature of focal adhesion organization. Thus, in addition to local development of traction force, microtubule targeting to focal adhesions should somehow weaken the adhesion strength making the adhesions prone to detachment by a transient local increase in actomyosin contractile force.

What are the microtubule-associated factors which could increase the probability of focal adhesion sliding upon transient local activation of contractility? One possibility is that KANK1 itself could somehow weaken the integrin adhesions and facilitate their sliding as shown in experiments by by Fassler’s group^17^. However, this contradicts to our observation that in cells lacking microtubules, recruitment of KANK to the focal adhesions does not disrupt them. Thus, the focal adhesion sliding induced by KANK overexpression is a microtubule dependent phenomenon and cannot be explained solely by an increase of KANK level in focal adhesions. Overexpression of KANK1 also did not reduce podosome numbers^16^ which is inconsistent with the idea of KANK-mediated weakening of talin-actin interaction^17^. Another possible reason for focal adhesion weakening immediately after microtubule attraction induced by OptoKANK activation could be the transient depletion of GEF-H1 in the proximity of the focal adhesion as predicted by our model. This could result in decrease of myosin activity, resulting in a reduction of pulling force experienced by focal adhesion and consequently partial focal adhesion disassembly. However, we did not detect any drop of traction force upon OptoKANK activation that preceded the traction force augmentation. In addition, the OptoKANK activation induced focal adhesion sliding also in cells treated with CNO3 that presumably have a high level of Rho activity. In such cells, depletion of GEF-H1 by microtubules recruited to focal adhesions will barely affect Rho activity level and reduces contractile forces experienced by the focal adhesions. Thus, the most plausible scenario is the delivery of focal adhesion weakening factors by microtubules.

To elucidate the process of microtubule driven focal adhesion disassembly we conducted an extensive screen revealing the involvement of key molecular players in this process. In the course of these studies, we did not find evidence of involvement specific proteolysis or endocytosis in the process of microtubule-driven focal adhesion disassembly. Indeed, inhibitors of calpain, matrix metalloproteases (MMP) and dynamin, did not interfere with focal adhesion disassembly induced by OptoKANK activation. Several other molecular players suggested in the literature were also not confirmed in our experimental system. In part, this discrepancy could be attributed to the difference between experimental systems: local targeting of microtubules to focal adhesions used in our study versus global microtubule outgrowth after nocodazole washout used in majority of previous studies.

We identified several proteins for which knockdown or/and inhibition efficiently prevented the microtubule-driven focal adhesion disassembly after OptoKANK activation. Besides the Rho activation axis (GEF-H1, ROCK) we found that kinases FAK and PAK, microtubule motor kinesin-1, tubulin acetylase αTAT1 and microtubule-associated activator of actin polymerization APC, are needed for the focal adhesion disassembly. The involvement of FAK, PAK and kinesin-1 is consistent with the previous publications^25, 33^ .

To clarify whether FAK, APC, and Kinesin-1 participate in local microtubule-driven activation of Rho and actomyosin contractility similar to GEF-H1, we investigated whether pharmacological activation of Rho during illumination of focal adhesions can overcome the effects of depletion or inhibition of these proteins. We discovered that suppression of PAK and GEF-H1 can be abolished by activation of Rho. This suggests that other players have different functions and, in particular, could participate in the microtubule-dependent focal adhesions weakening preceding their sliding induced by the burst of contractility. While a plausible function of kinesin-1 is a transportation of hypothetical factors weakening the focal adhesions along microtubules, the role of APC, FAK and αTAT1, in the process of focal adhesion weakening triggered by OptoKANK activation requires further studies. Of note, the identified players can work synergistically, as kinesin-1 is known to be involved in the transport of APC along microtubules^44^, while microtubule acetylation by αTAT1 can affect association of GEF-H1 with microtubules^38^.

While molecular details of microtubule-driven focal adhesion disassembly remain to be elucidated, our study provided strong evidence in favor of a new model of this process. We have shown that the key event induced by microtubule targeting focal adhesion is local GEF-H1 dependent activation of myosin filament formation and actomyosin contractility near the proximal end of the focal adhesion. This local contraction triggers the focal adhesion sliding which in turn results in its disassembly. Future studies will clarify how this mechanism functions during 2D and 3D cell migration and determine if this mechanism applies to the interaction of microtubules with other types of integrin adhesions like podosomes and fibrillar adhesions.

## Materials and methods

### Cell culture and cell transfection procedures

The HT1080 human fibrosarcoma cell line was obtained from the American Type Culture Collection and cultured in MEM supplemented with 10% heat-inactivated FBS, non-essential amino acids and sodium pyruvate (Sigma-Aldrich), in an incubator at 37 °C and 5% CO_2_. FAK^-/-^ MEF cells were a gift from P. Kanchanawong (Mechanobiology Institute, Singapore) and were cultured in DMEM supplemented with 10% heat-inactivated FBS, non-essential amino acids and sodium pyruvate (Sigma-Aldrich) and penicillin–streptomycin (Thermo Fisher Scientific), in an incubator at 37 °C and 5% CO_2_.

Cells were transiently transfected with the expression vector plasmids using electroporation (Neon Transfection System, Life Technologies) in accordance with the manufacturer’s instructions. Specifically, one pulse of 950 V of 50 ms was used for HT1080 cells, one pulse of 1350 V of 20 ms was used for the FAK^-/-^ MEF cells. For siRNA-mediated knockdown, HT1080 cells were transfected at the following concentrations: 100 nM for GEF-H1 (Dharmacon, ON-TARGETplus siRNA, cat. no. J-009883-09-0002), 100 nM for αTAT1 (ThermoFisher siRNA cat. no. AM16708), 100 nM for APC (Dharmacon, cat. no. L-003869-00-0005), 100 nM for BNIP2 (gift from Low B.C.), and 100 nM for ARP2. For control experiments, cells were transfected with non-targeting pool siRNA (ON-TARGETplus, Dharmacon cat. no. D-001810-10) at a concentration similar to that of the gene-targeted siRNAs. Cells were transfected using Lipofectamine RNAiMAX (Invitrogen) according to the manufacturer’s instructions.

### Plasmids

For generation of iLID-based optogenetic constructs of KANK (OptoKANK), the KN domain of KANK1 corresponding to amino acid residues 1–68 was fused with the mApple fluorescence tag and LOV2ssrrA domain (mApple–KANK-KN–LOV2ssrA)^26^. The rest of KANK1, ΔKN, corresponding to amino acid residues 69–1352, was fused with the mEmerald fluorescence tag and the SspB domain (SspB– ΔKN–mEmerald)^26^. The mApple–KN–LOV2ssrA and SspB–ΔKN–mEmerald constructs were cloned by Epoch Life Science. For generation of ITGB3-mIFP and mIFP-Talin1 constructs, the GFP fluorescent tags of ITGB3-GFP (gift from Jonathan Jones – Addgene plasmid #26653) and GFP-Talin1 (gift from Anna Huttenlocher – Addgene plasmid #26724) were swapped out with the mIFP fluorescent tag of the mIFP-Vinculin (kindly provided by Dr. Michael W. Davidson, Florida State University, FL, USA). Constructs were cloned by Epoch Life Science.

### Live cell observation

Pharmacological treatments were performed using the following concentrations of inhibitors or activators: 1 μM for Nocodazole (Sigma-Aldrich), 0.4 μM for Y-27632 dihydrochloride (Sigma-Aldrich), 20 mM for Acetyl-Calpastatin (Tocris, Cat. No. 2950), Monastrol (Merk, cat. No. 254753-54-3), 80 µM for Dynasore (Abcam Cat. No. 120192), Batimastat (Tocris Cat. No. 2961), Tubacin (Cat. No. 3402), MAP4K4-IN-3 (MedChemExpress, Cat. No.: HY-125012), 1 µM for FRAX1036 (MedChemExpress Cat. No. HY-19538), 50 µM for Kinesore (Cat. No. 6664), 5 µM for PF-57328 (Tocris, Cat. No. 3239), and 1 μg ml^-1^ for Rho activator II (CNO3, Cytoskeleton). Cells were treated at least 2h at 37 °C and 5% CO2, except when specified in the legend, before OptoKANK-based live imaging. For nocodazole-washout experiments, transfected HT1080 cells were plated on 35-mm dish from IBIDI overnight. One hour before imaging, nocodazole in fresh MEM medium with 10% FBS was added to the cells. The 35-mm dishes were mounted in a perfusion chamber (CM-B25-1, Chamlide CMB chamber). Nocodazole was washed out using fresh MEM medium with FBS 2 hours later. For the contractility rescued experiments, cells were first depleted 24h or treated with inhibitors for 2 hours before addition of CNO3 at 1 μg mg.ml^-1^. Cells were then imaged using microscope 1h after addition of CNO3 without changing the medium.

### Fluorescence microscopy

OptoKANK-based live experiments were performed using total internal reflection fluorescence (TIRF) microscopy (Olympus IX81, Zero Drift Focus Compensator, Dual camera Hamamatsu ORCA-Fusion BT, Objective x100) or using confocal microscopy (Yokogawa CSU-W1, Nikon TiE, 2x Photometrics Prime 95b CMOS camera, Objective X63) using Metamorph software. For OptoKANK activation, the 488 nm wavelength laser was set up at 1% in Fluorescence Loss in Photobleaching (FLIP) mode on the region of interested (ROI). Classical time-course experiments were performed at 5 sec intervals with continuous photoactivation (except every 5 sec corresponding to the acquisition steps), and 3 sec intervals for EB1 experiments.

### Immunoblotting

Cells were lysed directly in Laemmli buffer for Western blot (Tris–HCl pH 6.8 0.12 M, glycerol 10%, sodium dodecyl sulfate 5%, β-mercaptoethanol 2.5%, bromophenol blue 0.005%) and extracted proteins were separated by SDS–PAGE in 4–20% SDS–polyacrylamide gel (Thermo Fisher Scientific) and transferred to polyvinylidenedifluoride membranes (Bio-Rad) at 75 V for 2h.

Subsequently, the polyvinylidenedifluoride membranes were blocked for 1 h with 5% bovine serum albumin (BSA, Sigma-Aldrich), then incubated overnight at 4 °C with appropriate antibodies: anti-GEF-H1 (Cell Signaling, cat. no. 4145, dilution 1:1000); anti-α-tubulin (Sigma-Aldrich, cat. no. T6199, dilution 1:3000); anti-GAPDH (Santa Cruz Biotechnology, cat. no. sc-32233, dilution 1:3000); anti-APC (Thermo Fisher Scientific, cat. no. A5-35188, dilution 1:1000); anti-ARP2 (Cell Signaling, cat. no. #3128, dilution 1:1000); anti-Phospho-PAK1 (Ser199/204)/PAK2 (Ser192/197)4) (Cell Signaling, cat. no. #2605, dilution 1:2000); anti-acetyl-α-Tubulin (Lys40) (Cell Signaling, cat. no. #3971, 1:1000); anti-αTAT1 (Thermo Fisher Scientific, cat. no. PA5-112992, dilution 1:1000).

Subsequently, the membranes were washed three times (10 min each) and probed by incubation for 1 h with the appropriate secondary antibodies conjugated with horseradish peroxidase (Bio-Rad). The membranes were then washed three times (15 min each at room temperature), developed using PierceTM ECL western blotting substratum (Thermo Fisher Scientific) and imaged using a ChemiDoc imaging system (Bio-Rad).

### Micropatterning of adhesive islands

Fibronectin-patterned glass coverslips were microfabricated using the first steps of the glass technique described by Vignaud et al^48^. Briefly, glass coverslips (VWR) were plasma treated for 30 s and incubated for 30 min at room temperature with 0.1 mg.ml^-1^ poly-L-lysine-grafted-polyethylene glycol (pLL-PEG, SuSoS) diluted in HEPES (10 mM, pH 7.4, Sigma). After washing in deionized phosphate-buffered saline (dPBS, Life technologies), the pLL-PEG covered coverslip was placed with the polymer brush facing downwards onto the chrome side of a quartz photomask (Toppan) for photolithography treatment (5 min UV-light exposure, UVO Cleaner Jelight). Subsequently, the coverslip was removed from the mask and coated with 30 µg.ml^-1^ fibronectin (Sigma) diluted in dPBS for 30 min at RT.

### EB3 comet tracking and data analysis

Images of EB3-mIFP acquired from the TIRF microscope were exported as multidimensional TIFF files. EB3 comets from these raw unprocessed images were tracked automatically using the plusTipTracker software^49, 50^. To measure displacement, lifetime and velocities of EB3 comets, the following parameters were set in the program: search radius range, 4–15 pixel; minimum subtrack length, 3 frames; maximum gap length, 10 frames; maximum shrinkage factor, 0.8; maximum angle forward, 50; maximum angle backward, 10; and fluctuation radius, 2.5. To visualize comet tracks in individual growth cones, the plusTipSeeTracks function was used. MT dynamics parameters were compiled from multiple individual experiments.

### Traction force microscopy

PDMS micropillars were fabricated to form PDMS mold for micropillar array. After silanizing the surface of the PDMS pillars with Trichloro (1H,1H,2H,2H-perfluorooctyl) silane (Sigma, 448,931) overnight, new PDMS (DOWSIL 184 silicone elastomer, Dow Corning, MI, USA) was directly cast onto the surface of the micropillar to make a PDMS mold with holes. After degassing for 15 min, the mold was cured at 80 degrees for 2h. The PDMS mold was peeled off from the PDMS pillars, cut 1 cm square and placed on plastic dishes face up following a silanization of their surface with Trichloro (1H,1H,2H,2H-perfluorooctyl) silane overnight.

To fabricate the array of micropillar, whose refraction index is similar to that of the growth medium, a small drop of My-134 polymer (My Polymers Ltd., Israel) was put on the center of coverslip coated with 3-(Trimethoxyilyl)propyl methacrylate (sigma, 440,159) and then, the silanized PDMS mold covered the drop face down onto the coverslip with thin layer of My-134 polymer for 15–30 min. After degassing for 5–15 min to get rid of air bubbles inside the polymer, the assembly was placed in a cell culture dish, covered with fresh milli-Q water and cured under short wavelength UV radiation (UVO Cleaner 342A-220, Jelight Company Inc., USA) for 6 min. Then, the PDMS mold was carefully peeled off from the coverslip.

Top of My-134 pillars were coated with fluorescence-labelled fibronectin. Briefly, PDMS stamps were incubated with solution containing 30 μg/ml fibronectin and 2 μg/ml fibrinogen Alexa Fluor 647 conjugate solution (F35200, Invitrogen, USA) in Dulbecco’s phosphate-buffered saline (Sigma-Aldrich) at room temperature for 90 min. After washing with Milli-Q water and air-drying the surface, the PDMS stamp was put onto the top of My-134 pillars freshly exposed to UV-Ozone (UV Ozone ProCleaner Plus, BioForce Nanosciences). After 5 min of contact, the stamp was removed. Before cell plating, the My-134 pillars were incubated with 0.2% Pluronic F-127 (Sigma) for 1 h for blocking, followed by washing three times with Dulbecco’s phosphate-buffered saline. The pillars in the array were arranged in a triangular lattice with 4 μm center-center distance and the dimensions of pillars were d = 2.1 μm with h = 6 μm (k = 50 nN/μm). The traction forces by fluorescent-labelled My-134 pillars were calculated using a custom-build MATLAB program (version 2019a, MathWorks) as described previously^51^.

### Immunofluorescence microscopy

Cells were fixed with 3.7% PFA (Sigma) for 20 min, permeabilized with 0.1% Triton in PBS (Sigma), then washed with PBS, and blocked with a blocking solution (2.5% bovine serum albumin in PBS Tween 0.2%) for 1 h. Samples were incubated overnight at 4°C with primary antibody in blocking solution : anti-vinculin (Sigma-Aldrich, catalogue no. V9131, dilution 1:400) followed by three washes with PBS Tween 0.2%. The cells were then incubated with secondary antibody at room temperature for 1 h followed by three washes with PBS Tween 0.2%.

### Physical model

See supplemental methods.

### Statistical analyses

Statistical analyses were performed using GraphPad Prism software (GraphPad, version 9). Statistical significance was determined by the specific tests indicated in the corresponding figure legends.

## Supporting information

Model - supplemental methods

Supplemental movie 1 - Ctrl

supplemental movie 2 - Nocodazole

Supplemental movie 3 - washout

Supplemental movie 4 - EB3 comets

Supplemental movie 5 - Ctrl vs Dynasore

Supplemental movie 6 - MLC

Supplemental movie 7 - SiR Actin

Supplemental movie 8 - Ctrl and GEF-H1 KD

Supplemental movie 9 - GEF-H1 KD + CNO3

## Acknowledgements

We thank Christophe Guilluy for support and encouragements, A. Wong (MBI, Singapore) for expert help in paper editing, Diego Pitta de Araujo (MBI, Singapore) for artistic design of the figure 6, Bryant L. Doss for advices with the TFM experiments, Yukako Nishimura (Hokkaido University) for advices with the MY-134 pillars technology, M. Davidson fluorescence protein collection (The Florida State University, Tallahassee, USA), the SIMBA microscopy facility and nanofabrication core facility at the Mechanobiology Institute for technical help. The research is supported in part by the Singapore Ministry of Education Academic Research Fund Tier 2 (MOE Grant No: MOE2018-T2-2-138, awarded to A.D.B; MOE2019-T2-1-099 and MOE2019-T2-02-014; awarded to P.K.), and Tier 3 (MOE Grant No: MOE2016-T3-1-002 and MOET32021-0003; awarded to A.D.B), the National Research Foundation, Prime Minister’s Office, Singapore, and the Ministry of Education under the Research Centers of Excellence program through the Mechanobiology Institute, Singapore (ref no. R-714-006-006-271).

## Author contributions

J.A. and A.D.B conceived and designed the experiments. J.A. performed all experiments. S.B. and K.P. helped with the OptoKANK construct design, L.L, I.A., and B.C contributed to data analysis and design of the paper, A.M. created and analyzed the physical model and, J.A, A.M and A.D.B wrote the manuscript with input from all of the authors.

## Supplemental figures

**Supplemental figure 1.**
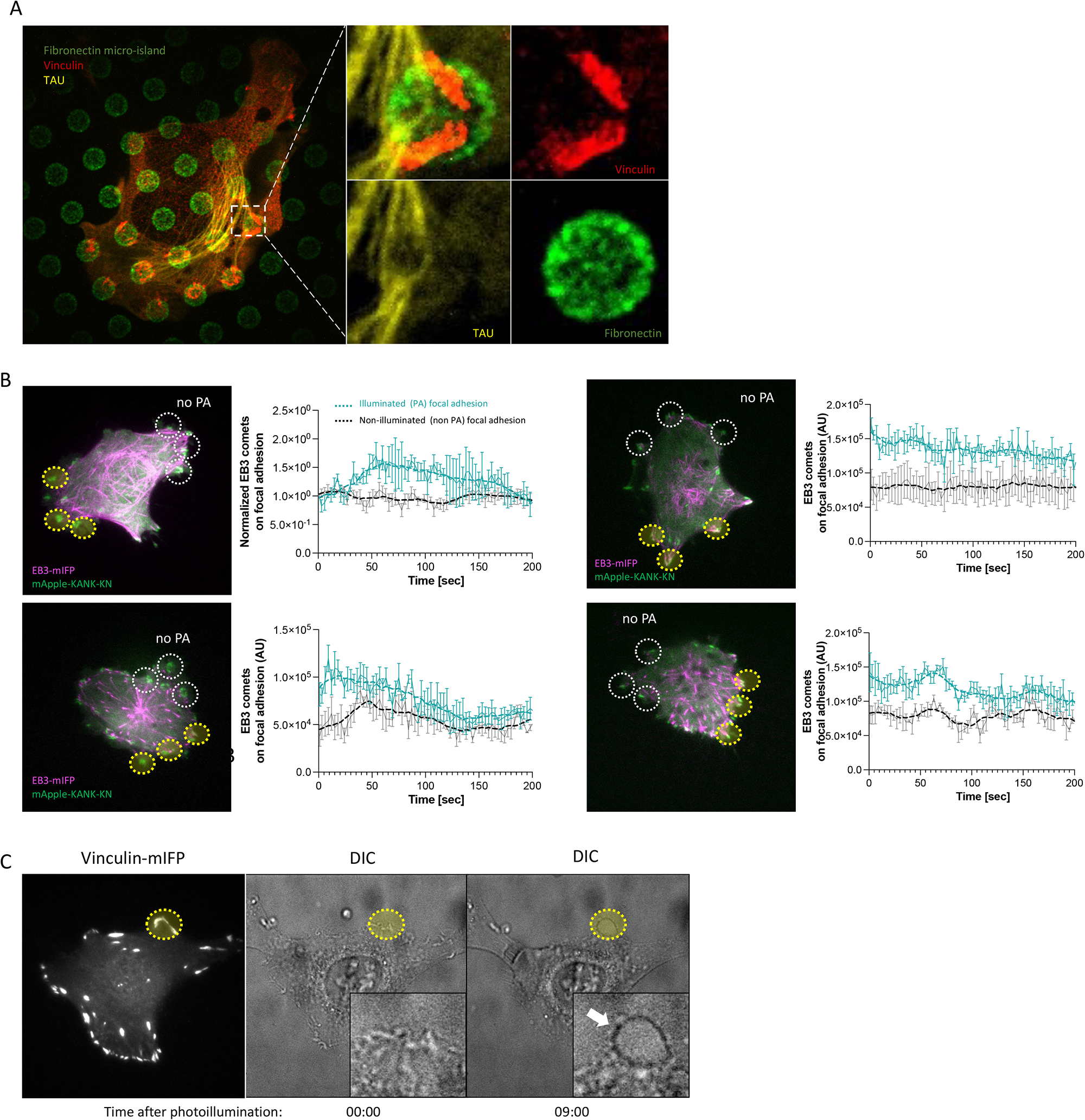
**(A)** Example of HT1080 cell transfected with TAU-mCherry, plated on microislands pattern labelled in green (Alexa 647) and stained for focal adhesions using anti-vinculin (red). Typically, the cell displays 2 to 4 focal adhesions par island where the microtubules visualized thanks to TAU protein (yellow) are passing by the focal adhesion. **(B)** Graphs show the integrated EB3 comet fluorescence after processing with U-Track2 for photoactivated and non-photoactivated focal adhesions over the time. (Data are presented as the mean of integrated EB3 fluorescence from 3 focal adhesions under each condition). **(C)** Representative images of Vinculin-mIFP-transfected HT1080 cells carrying the OptoKANK constructs after treatment with Kinesore. Images show the illuminated focal adhesion (yellow dotted circle) using vinculin-mIFP staining and the DIC images showing the bleb formation (see white arrow in the box) after 9 min of OptoKANK activation.

**Supplemental figure 2.**
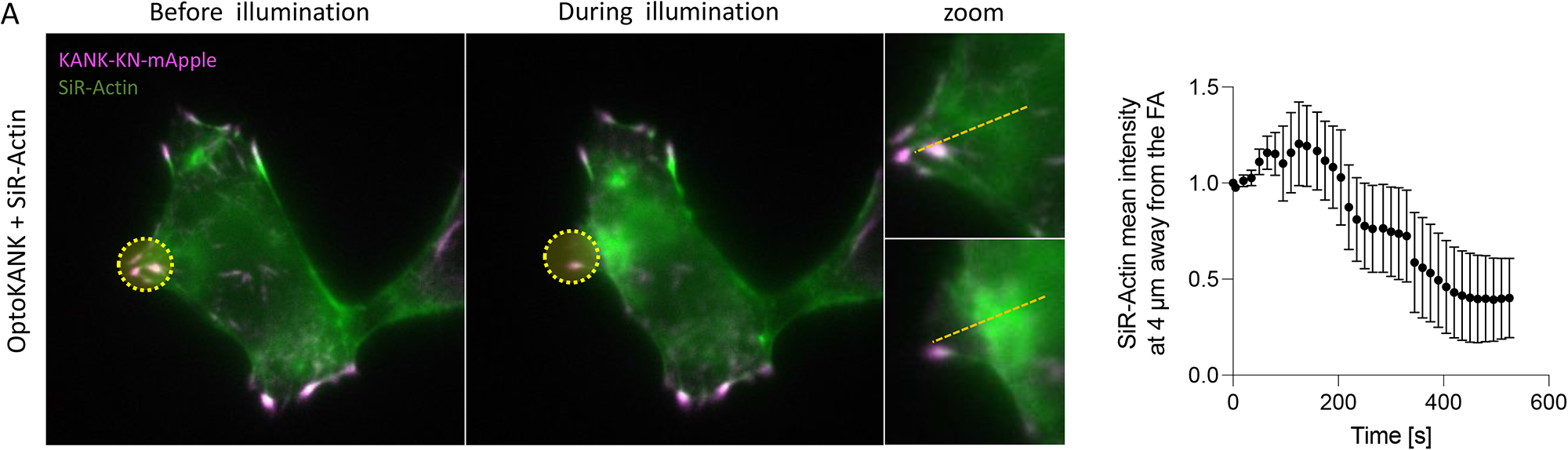
**(A)** Representative image of OptoKANK-transfected HT1080 cell before and during OptoKANK activation of the selected focal adhesion (yellow dotted line). SiR-Actin was used to assess the myosin-II dynamics upon OptoKANK activation. The line scan was used to measure the SiR-Actin intensity in the vicinity (4 µm away in centripetal direction) of the proximal end of the photoactivated focal adhesion shown. The graph shows the SiR-Actin mean intensity at this distance upon OptoKANK activation (Data are presented as mean ± s.e.m; n = 7).

**Supplemental figure 3.**
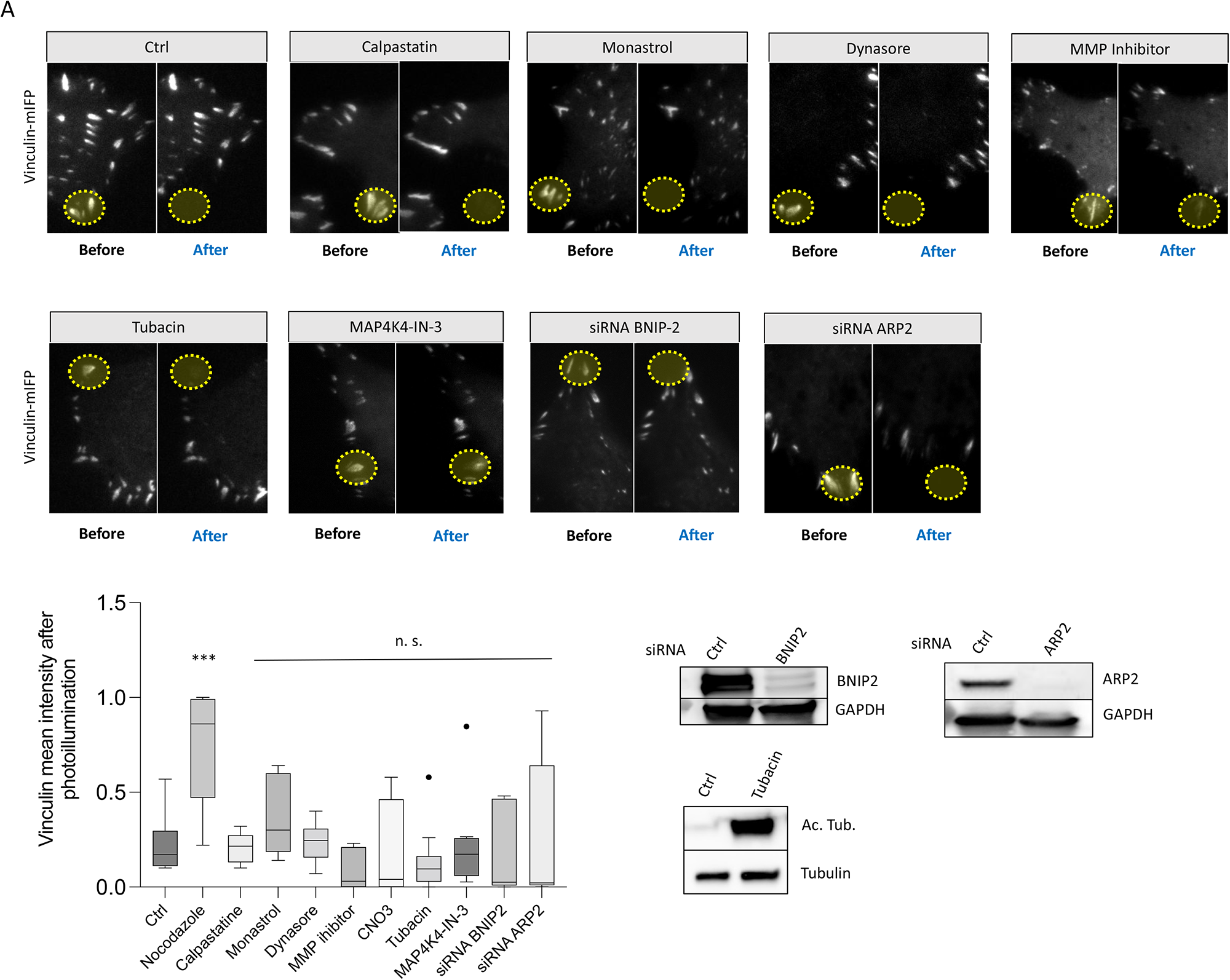
**(A)** Representative images of Vinculin-mIFP-transfected HT1080 cells carrying the OptoKANK constructs before and after blue light illumination of the focal adhesion (yellow dotted line) for control cells, cells treated with a calpain inhibitor, calpastatin, a kinesin-V inhibitor, monastrol, a dynamic inhibitor, dynasore, a MMP inhibitor, Batimastat, a MAP4K4 inhibitor, MAP4K4-IN-3, and depleted for BNIP2 and ARP2. Graph shows the normalized mean vinculin intensity after the illumination of cells treated as indicated (Data are presented as mean ± s.e.m; Ctrl, n =19; Nocodazole, n = 12; Calpastatin, n = 10; Monastrol, n = 8; Dynasore, n = 10; MMP inhibitor, n = 7; CNO3, n = 8; Tubacin, n = 12, MAP4-K4-IN3, n = 9; siRNA BNIP2, n = 16; siRNA AR2, n = 10). Immunoblots of BNIP2, ARP2, acetylated tubulin and GAPDH are shown in the black box.

**Supplemental Figure 4.**
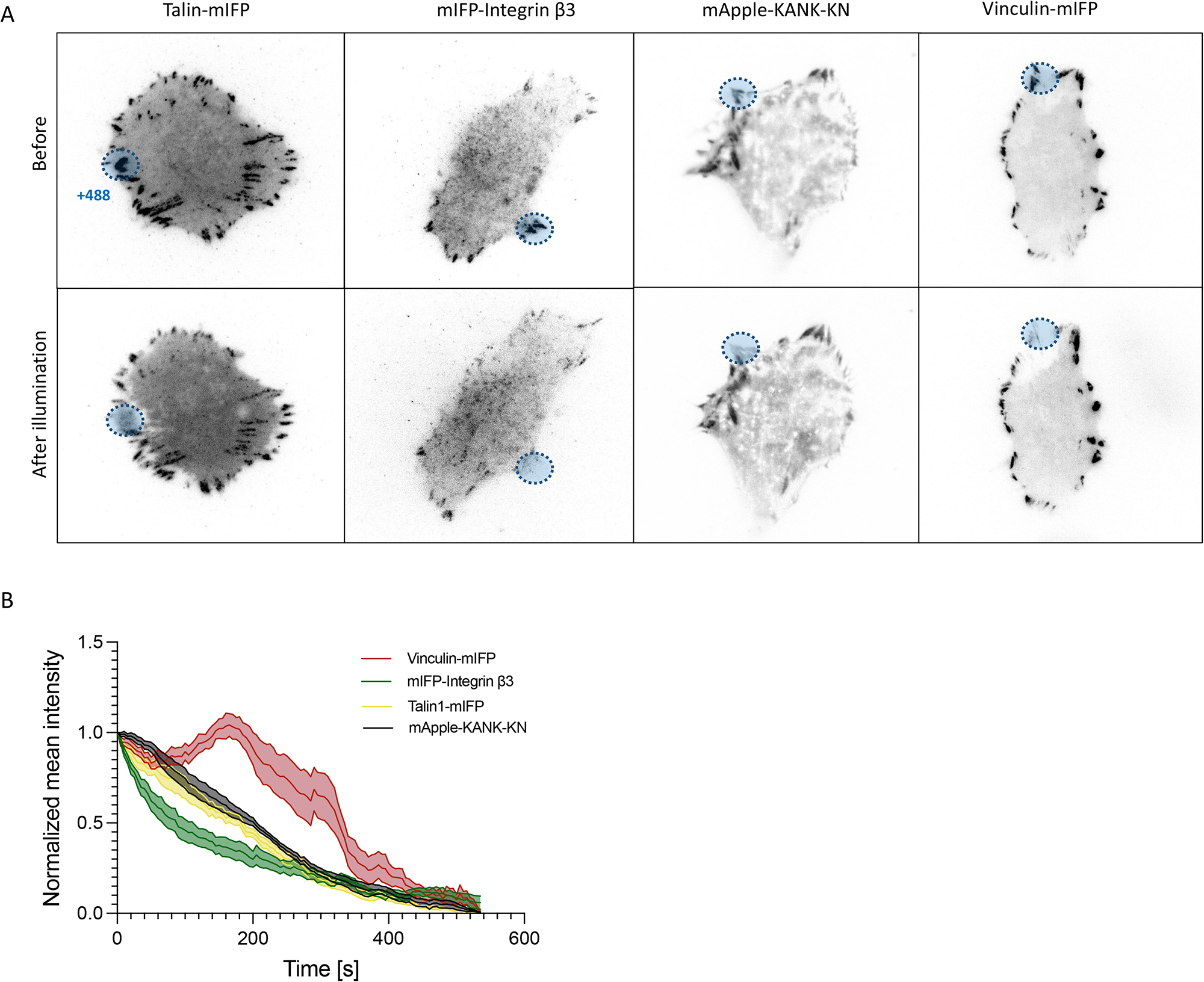
**(A)** Representative images of HT1080 cells carrying the OptoKANK constructs before and after blue light illumination of areas (blue dotted line) containing focal adhesion visualized using Talin-mIFP, mIFP-Integrin β3, mApple-KANK-KN and Vinculin-mIFP labeling. **(B)** Normalized mean intensity of Vinculin (vinculin-mIFP), Integrin β3 (mIFP-Integrin β3), Talin1 (Talin1-mIFP) and KN (mApple-KANK-KN) over the illuminated focal adhesion of cells carrying OptoKANK.

**Supplemental Figure 5.**
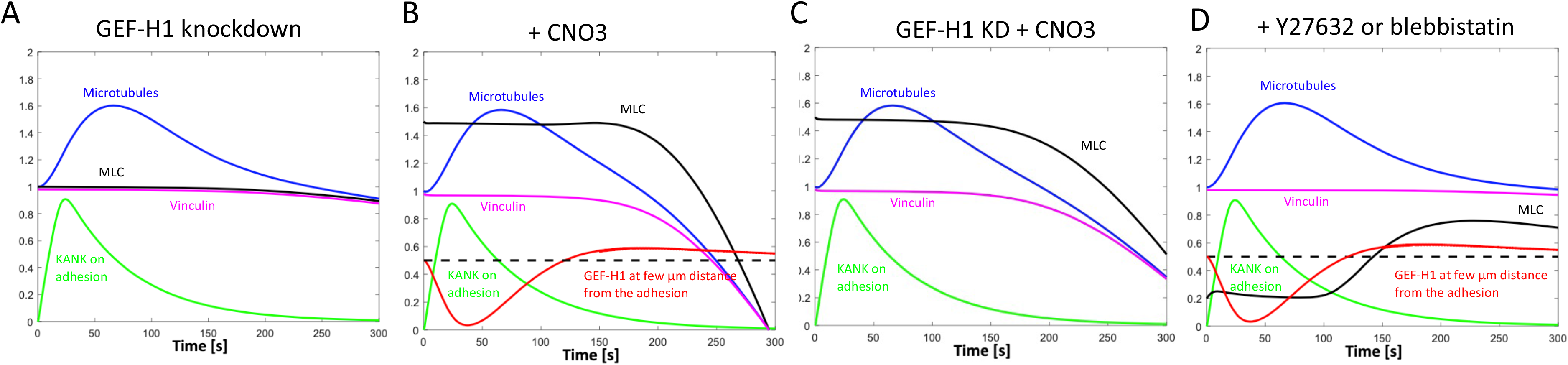
Model-predicted time series for microtubule, vinculin, KANK, GEF-H1 and myosin densities for the following simulated perturbations: GEF-H1 knockdown **(A)**, treatment with CNO3 **(B)**, GEF-H1 knockdown + CNO3 **(C)** and treatment with Y27632 or blebbistatin **(D)**. Model parameters used in the simulations are described in the Supplemental Text.

**Supplemental Figure 6.**
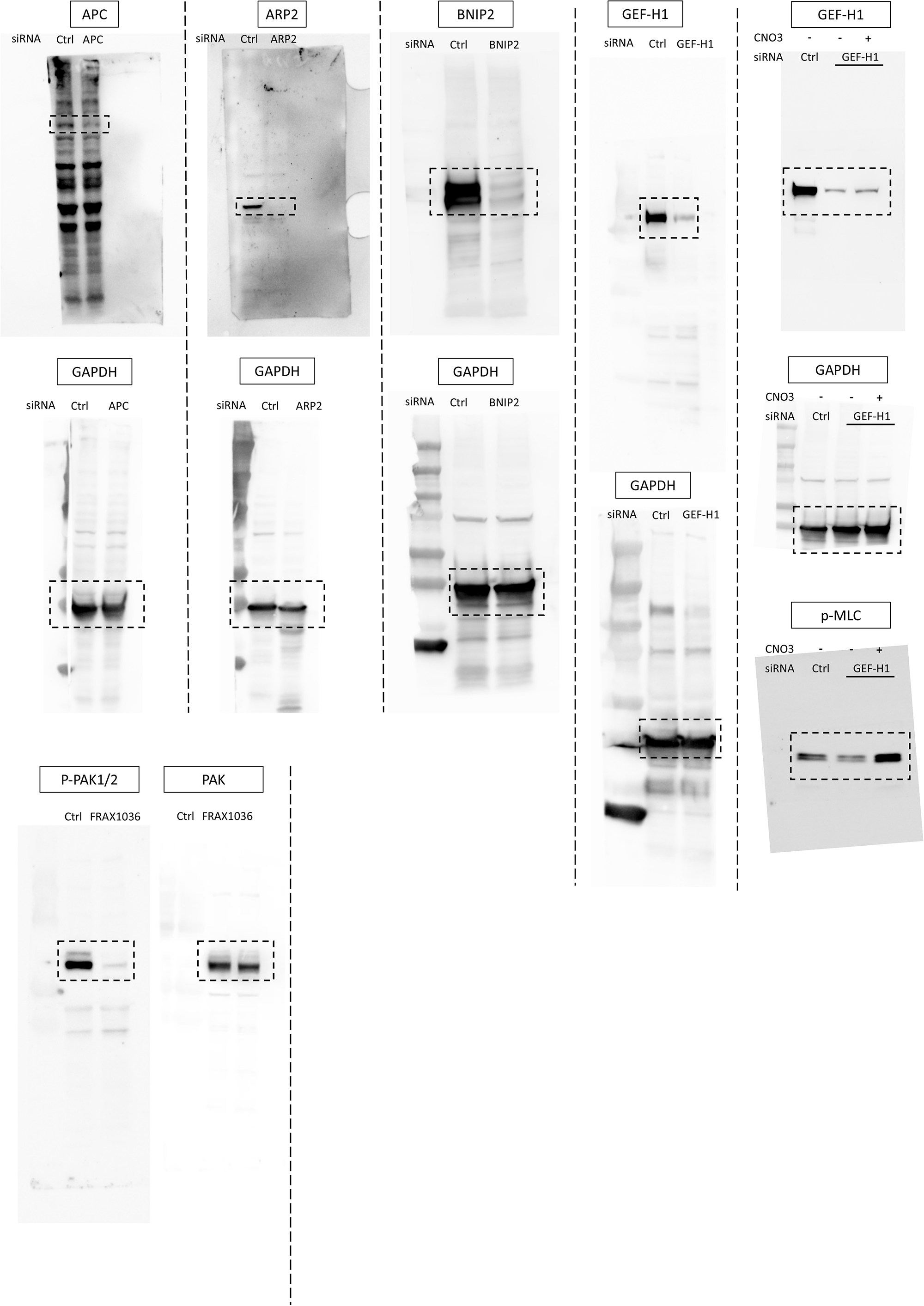
Uncropped images of western blots of HT1080 cells showed in Figure 4B and 4C, Figure 5A and supplemental figures 3A. The boxed areas (black frames) show the cropped images.

## Supplemental movie legends

### Supplemental movie 1

**OptoKANK activation promotes focal adhesion sliding and disassembly**

HT1080 cell transfected with OptoKANK (KN + ΔKN) and vinculin-mIFP was illuminated (488 nm) over the focal adhesion; approximate area of illumination is demarcated by the blue circle. Typical movie in which the focal adhesions labeled by vinculin-mIFP slide and disassemble upon OptoKANK activation. Acquisition rate is 1 frame/5 sec and display rate is 30 frames/sec.

### Supplemental movie 2

**Nodazole treatment prevents focal adhesion disassembly upon OptoKANK activation**

HT1080 cell transfected with OptoKANK (KN + ΔKN) and vinculin-mIFP was illuminated (488 nm) over the focal adhesion; approximate area of illumination is demarcated by the blue circle. The cell was treated with nocodazole (1µM) for 3 hours prior illumination and imaging of the focal adhesion labeled by vinculin-mIFP. Note that in the nocodazole treated cell, the focal adhesion remains unchanged after OptoKANK activation. Acquisition rate is 1 frame/5 sec and display rate is 30 frames/sec.

### Supplemental movie 3

**Nocodazole washout rescues focal adhesion sliding and disassembly upon OptoKANK activation**

HT1080 cell transfected with OptoKANK (KN + ΔKN) and vinculin-mIFP was illuminated (488 nm) over the focal adhesion (in the blue circle). The cell was treated with nocodazole (1µM) for 3 hours followed by 2 hours washout prior illumination and imaging of the focal adhesion labeled by vinculin-mIFP. A typical movie in which the focal adhesion slides and disassembles upon OptoKANK activation. Acquisition rate is 1 frame/5 sec and display rate is 30 frames/sec.

### Supplemental movie 4

**OptoKANK activation promotes microtubule tips targeting to focal adhesion**

HT1080 cell transfected with OptoKANK (KN + ΔKN) and EB3-mIFP was illuminated (488 nm) over the focal adhesions (in the blue circles) and the EB3 comets (left panel) were tracked automatically using the plusTipTracker software^49, 50^ (right panel). Non illuminated focal adhesion areas (white circles on left panel and dark circles on right panel) were used as control. Acquisition rate is 1 frame/3 sec and display rate is 20 frames/sec.

### Supplemental movie 5

OptoKANK activation results in myosin-II filaments accumulation in the vicinity of the illuminated focal adhesion proximal end.

HT1080 cell transfected with OptoKANK (KN + ΔKN) and MLC-mIFP was illuminated (488 nm) over the focal adhesion (yellow circle). The position of focal adhesion is indicated in the first frame by KN-mApple fluorescence. Accumulation of myosin-II filaments based on the MLC-mIFP intensity appears in centripetal direction from focal adhesions following the onset of illumination (see white arrow). Acquisition rate is 1 frame/5 sec and display rate is 10 frames/sec.

### Supplemental movie 6

**OptoKANK activation results in actin accumulation in the vicinity of the illuminated focal adhesion proximal end.**

HT1080 cell transfected with OptoKANK (KN + ΔKN) and pre-incubated with SiR-actin for 1h was illuminated (488 nm) over the focal adhesion (yellow circle). The focal adhesion is visualized using KN-mApple fluorescence. Accumulation of actin filaments visualized by the SiR-actin labeling appears in centripetal direction from focal adhesions following the onset of illumination. Acquisition rate is 1 frame/5 sec and display rate is 30 frames/sec.

### Supplemental movie 7

**GEF-H1 knockdown prevents focal adhesion disassembly upon OptoKANK activation**

HT1080 cells transfected with OptoKANK (KN + ΔKN) and vinculin-mIFP was illuminated (488 nm) over the focal adhesion (blue circle) in control (left panel) or after GEF-H1 knockdown (right panel). While the focal adhesion slides and disassembles upon OptoKANK activation in ctrl, depletion of GEF-H1 typically prevents the sliding and disassembly of focal adhesion. Acquisition rate is 1 frame/5 sec and display rate is 20 frames/sec.

### Supplemental movie 8

**Rescuing the GEF-H1 knockdown effect on focal adhesion disassembly upon OptoKANK activation by activation of RhoA with CNO3**

GEF-H1-knocked down HT1080 cell transfected with OptoKANK (KN + ΔKN) and vinculin-mIFP was illuminated (488 nm) over the focal adhesion (yellow circle). CNO3 (1 µg/ml) was added 2h prior the onset of illumination. Upon OptoKANK activation, the focal adhesion slides and disassembles in spite of GEF-H1 depletion. Acquisition rate is 1 frame/5 sec and display rate is 20 frames/sec.

### Supplemental movie 9

**Inhibition of endocytosis results in local membrane bleb formation upon OptoKANK activation but does not protect focal adhesion**

HT1080 cells transfected with OptoKANK (KN + ΔKN) and vinculin-mIFP was illuminated (488 nm) over the focal adhesion (yellow circle) in control (left panel) or in the presence of the endocytosis inhibitor, dynasore (right panel). OptoKANK activation of cell treated with dynasore results in formation of membrane bleb at the illuminated sites (phase contrast) and induces sliding and disassembly of focal adhesion as visualized by vinculin-mIFP (red). Acquisition rate is 1 frame/5 sec and display rate is 20 frames/sec.

