## Supplementary material for "Focal adhesions are controlled by microtubules through local contractility regulation": Model - supplemental methods

The conceptual model is shown schematically in Figure 6. Model variables and parameters are listed in Supplemental Tables 1 and 2, respectively. Below, we discuss the dimensions and parameter values in detail. We discuss the model assumptions along with describing mathematical terms in the model equations. The first three model equations introduce the dynamics of the number of microtubules (MTs) on the focal adhesion (FA), number of KANK molecules on the FA, and active myosin density proximal to the FA, respectively:

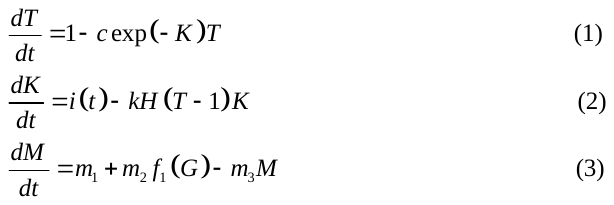

Here,
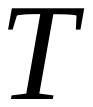
 is the number of MT plus ends on the FA,
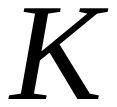
is the number of KANK molecules on the FA, and
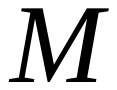
is the active myosin density proximal to the FA. The model does not describe explicit spatial molecular distributions, and so we only follow the temporal dynamics. Furthermore, even though MT and KANK numbers are not large, we approximate these numbers with continuous variables and from here on calling them ‘densities’ (these densities can be thought of as the respective numbers divided by the FA area). Also, we neglect stochasticity of these numbers and consider deterministic continuous model. Lastly, all densities in the model are non-dimensional, measured in units of characteristic observed scales. In the conceptual model, in the absence of the data on density dependence of relevant chemical rates, the dimensional numbers for the variables are not crucial.

Left-hand-sides of Equations 1-3 are the rates of change of the respective variables; we now turn to description of the right-hand-sides’ terms. The first term in Equation 1 describes arrival of the polymerizing growing MTs at a constant rate at the FA. This non-dimensional rate is chosen to be equal to unity because of the scaling discussed in the previous paragraph. The constancy of this rate is reasonable based on our observations suggesting that near the cell margins, the overall MT number and average dynamic instability parameters away from but proximal to adhesions do not exhibit noticeable variations. The second term is responsible for MTs leaving the FA with the rate equal to
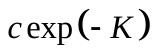
, where
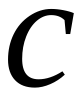
is the inverse pause time for MT prior to MT disassembly, before KANK activation. This parameter value can be estimated based on the observation of the MT pauses on the order of tens of seconds on the FAs (Azoitei et al 2019). The factor
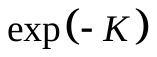
is responsible for the increase of the MT pause on the FA (and so the rate of leaving decrease) with growing KANK number (Bouchet2016). The exact functional form of this dependence is unknown, and the exponential form we use is a reasonable assumption. Implicitly, we use another assumption – that this rate decreases a few-fold when KANK is fully loaded onto the FA upon the illumination – which is based on the observed 1.5-fold increase of the MT number on the FA after the illumination. Note also that in principle there could be an effect of the MT catastrophe being promoted by the attachment of microtubule to the FA (Efimov and Kaverina 2009), but this effect can be effectively accounted for by the constant parameter
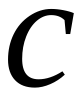
.

In Equation 2, the first term describes loading of KANK to the FA upon the illumination, as the following function of time:

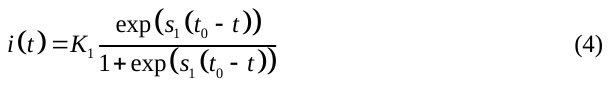

This function is a smoothed step function equal to constant,
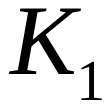
, before time
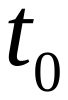
, and zero after this time. Parameter
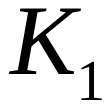
(Supplemental Table 2) is chosen so that by the end of the loading, KANK density reaches a characteristic unit; for
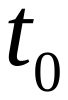
(Supplemental Table 2), we use the observed time of 20-30 seconds; smoothing parameter
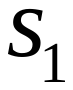
(Supplemental Table 2) is chosen to account for a sharp step-function-like transition. The second term in this equation accounts for the observed dissociation of KANK molecules from the FA on the scale of tens of seconds (constant parameter
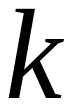
(Supplemental Table 2)). We also assume that MTs bring with them an adhesion-weakening molecule (Yue et al 2014), which triggers dissociation of integrins and talins, and because KANK binds to talin, of KANK molecules. This effect is accounted for by the factor
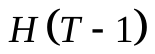
, where
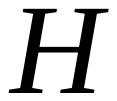
is Heaviside function equal to zero when the MT density is less or equal to unity, and to one otherwise. Thus, we assume that this effect has a threshold character. Another possibility is that KANK proteins themselves, without the MTs, diminish the talin-actomyosin linkage, which curbs force transmission across integrins, leading to reduced integrin–ligand bond strength, slippage between integrin and ligand, and sliding (Sun et al 2016). In that case, the factor
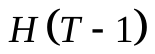
must be absent from Equation 2, and instead the dissociation rate simply equals constant
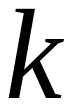
. The model’s results compare to the observations equally well with both choices.

In Equation 3, the first term is the constant (
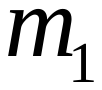
 (Supplemental Table 2)) basal, GEF-H1-independent activation rate of myosin, and the third term is the respective deactivation rate (which is assumed to be the first order chemical reaction with rate
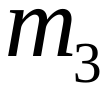
 (Supplemental Table 2)). The constant
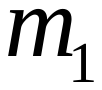
is chosen to bring the basal active myosin density to unity; the constant
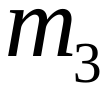
is chosen based on MLC phosphorylation cycle on the order of tens of seconds (Amano et al 1996). Note that in order to not overwhelm the model with molecular complexity, we do not explicitly describe the intermediate steps, like action of RhoGTPases, activation of ROCK, which are relatively fast, on the order of seconds (Bolado-Carrancio et al 2020) and can be lumped together with MLC phosphorylation rate. We also do not explicitly describe myosin clusters’ assembly. In the model, the myosin density is assumed to be proportional to the effective traction force and is a harbinger of the observed pMLC density.

The second term in Equation 3 is the GEF-H1-dependent rate of myosin activation. Rate
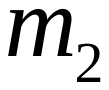
(Supplemental Table 2) characterizes the relative effect of GEF-H1-dependent compared to GEF-H1-independent rate. This parameter is unknown and chosen to fit the data. Function
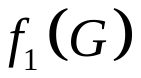
accounts for assumed threshold character of the GEF-H1 action:

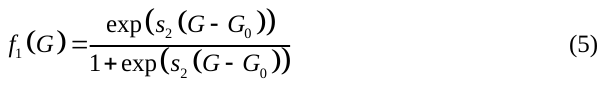

Here
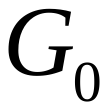
is the characteristic threshold density of activated GEF-H1 in the cytoplasm in the vicinity of the assembling actomyosin array proximal to the FA. Below this threshold, there is no effect, above the threshold there is a saturated effect. The threshold value is unknown and is chosen to fit the data (Supplemental Table 2). Parameter
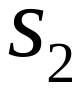
(Supplemental Table 2) characterizes the sharpness of the threshold effect; it is unknown from the experiment and is chosen to fit the data.

To close the system of model equations, we need GEF-H1 density as a function of time. The following equation provides such density at distance
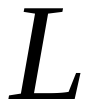
from the FA:

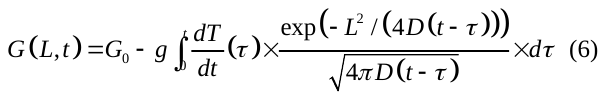

This formula is based on the following assumptions and estimates. First, MTs are effectively a sink for GEF-H1 molecules (Krendel2002). Thus, we assume that any time a MT arrives at the FA, a certain number of GEF-H1 molecules, equal to
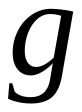
, is locally absorbed from the cytoplasm. The number of MTs arriving per time
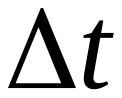
is
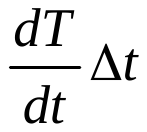
, and so
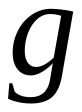

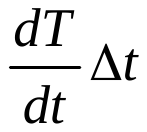
molecules are absorbed during this time interval. This expression assumes that the rate of absorption is independent of the local GEF-H1 concentration in the cytoplasm, so the limiting factor in the absorption is the local MT length/number and not GEF-H1 diffusion. Value of parameter
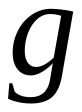
(Supplemental Table 2) is unknown and chosen to fit the data. Similarly, if MTs are leaving the FA but not arriving,
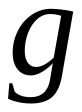

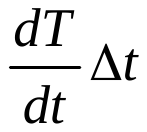
molecules are released during this time interval, assuming that GEF-H1 molecules are released from tubulin instantly upon the MT disassembly, so there is then a local source of GEF-H1 molecules on the FA. Neglecting local sinks and sources elsewhere in the vicinity, the spatially explicit concentration of GEF-H1 molecules in the cytoplasm can be described by the following reaction-diffusion equation:

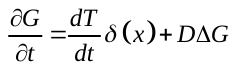

Here
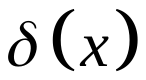
is the delta-function responsible for the location of the source/sink at the FA location,
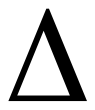
is the Laplacian operator, and
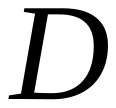
is the diffusion coefficient of GEF-H1 in the cytoplasm. We use a one-dimensional approximation of the geometry where
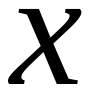
becomes the distance inward from the cell margin. In this approximation, the exact analytical solution for this reaction-diffusion equation is Equation 6 (Strauss 2007), where now
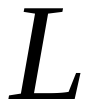
is the distance between the FA and proximal actomyosin array. Parameter
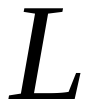
(Supplemental Table 2), based on our data is several microns. Value of parameter
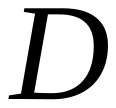
 (Supplemental Table 2), on the order of square microns per second, can be indirectly estimated from the data reported in (Azoitei et al 2019). To not overwhelm the model with molecular complexity, we do not explicitly include the step of activation of the released GEF-H1 molecules (Azoitei et al 2019), however, such process will not qualitatively change the conclusions from the model. One possible variant of the model is that we can in principle include the acetylation of MTs at the FA, and subsequent release of GEF-H1 from the MTs independent of the MT disassembly, since GEF-H1 has a low affinity to the acetylated MTs (Seetharaman et al 2022). Such variant of the model will lead to the same conclusions as our basic model.

Lastly, the mechanics of the FA slippage is accounted for in the model as follows. We assume that the pulling force applied to the FA, and therefore the traction force is directly proportional to the myosin density in the model. Next, we assume that the FA is ‘gripping’ if the ratio of the pulling force to adhesive strength is below a threshold, and slipping, if this ratio is above the threshold. We assume that the adhesive strength is proportional to the KANK density on the FA (not that KANK directly contributes to the adhesive strength, but rather that its adhesive molecular partners do, and their numbers are proportional to the KANK numbers), and that there is some basal, KANK-independent part of the strength. Mathematically, this translates into the assumption that the adhesive strength is proportional to the expression
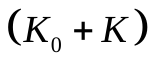
, where
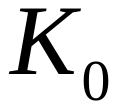
is a constant parameter. Then, the ratio of the pulling force to adhesive strength is proportional to the expression
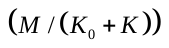
, and we define the velocity of slippage as follows:

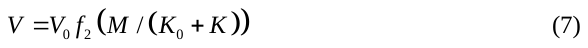

Here
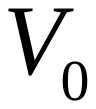
is the characteristic magnitude of the slippage velocity (Supplemental Table 2) that can be estimated from our data. Threshold function
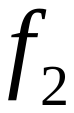
is defined as follows:

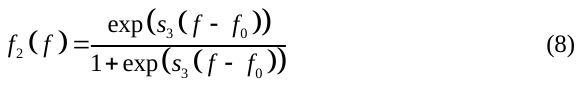

Here
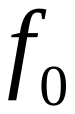
is the threshold force to adhesion strength ratio and
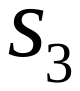
is the sharpness of the threshold parameter. The values of parameters
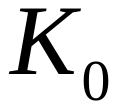
,
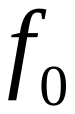
and
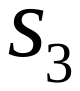
are unknown and chosen to fit the data (the results are insensitive to the value of
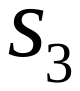
). The shift of the FA due to the slipping is given by the formula:

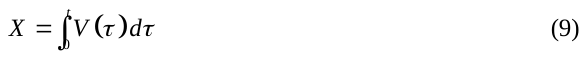

There is a gradual physical slipping of MTs, KANK and myosin, together with the slipping FA, and so we assume that the measured MT, KANK and myosin densities on the area of the initial adhesion scale with the initial adhesion length, on the order of one micron, minus the shift. Therefore, when the shift becomes equal to the initial adhesion length, the measured densities approach zero. Thus, as a result, we plot expressions
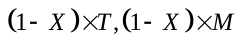
for the measured MT and myosin density. KANK density decreases so significantly by the time the slippage starts, that factor
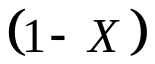
does not affect the result. Finally, we assume that the measured vinculin density is effectively a marker for the slipping adhesion area and approximate the measured vinculin density with expression
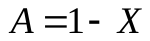
.

**Results:** The model equations are integrated by using standard Euler numerical scheme. The results are shown in Figure 6F. Here is the qualitative explanation for the predicted molecular densities as functions of time. KANK is rapidly loaded on the FA upon illumination, and then, as a part of the gradual, MT-induced weakening of the FA, starts to decrease exponentially. The transient accumulation of KANK causes longer pauses of the MTs on the FA, so the MT density on the FA increases at first, up to ~ 1.5-fold its value before the illumination, but after ~ 60 sec starts to decrease because the diminishing KANK number leads to shorter MT pauses on the FA.

Up to ~ 60 sec after the illumination, MTs arriving to the FA locally sequester GEF-H1 molecules, sharply depleting GEF-H1 density near the FA, but after that the decreasing MT number leads to releasing the accumulated GEF-H1 molecules. If the GEF-H1 dynamics was strictly local, then GEF-H1 molecules would be first captured by the MTs, then released in the same place, with no net gain. Crucially, because of diffusion, and because the sink on the increasing MT number occurs earlier than the source from the decreasing MT number, GEF-H1 molecules initially diffuse closer to the FA, down the gradient created by the sink. And then, when GEF-H1 is released by disassembling MTs, the net GEF-H1 concentration, after a short delay increases. Effectively, the initial MT increase, counter-intuitively, helps by ‘soaking’ GEF-H1 into the FA vicinity, then releasing the increased amounts of GEF-H1. Because the diffusion-reaction process introduces delays, GEF-H1 concentration becomes greater than the baseline after ~ 120 sec. This leads to a significant additional activation of myosin and traction force increase by ~ 150 sec. By that time, the adhesion is significantly weakened, the force to adhesive strength ratio exceeds the threshold, and the combined process of adhesion weakening and of local GEF-H1/RhoA/ROCK dependent activation of contractility leads to the slippage. These predictions compare well with the data (Figure 3).

The model also correctly accounts for the results of several perturbations experiments (Supplemental Figure).

*GEF-H1 knockdown*: we modeled this case by changing a single parameter – reducing the GEF-H1-dependent myosin activation rate,
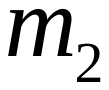
, to zero. As seen from the Supplemental Figure, in this case the illumination did not result in increase of traction force and sliding. Note a subtle shift of the FA in the end due to the weakening of adhesion, which leads to very slight slippage, as there is still basal myosin pulling action.

*Rho activator by CNO3*: We model this experiment by a 1.5-fold increase of the basal myosin activation rate
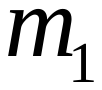
and by drastic decrease (Supplemental Table 2) of the GEF-H1-dependent myosin activation rate,
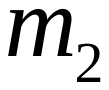
, assuming that most myosin is activated constitutively, minimizing additional effect of GEF-H1. The model predicts, correctly, that the FA sliding takes place (Supplemental Figure). The explanation is straightforward: before illumination, there is an elevated pulling on the FA, but it is still below the threshold that causes sliding. After the illumination, the MTs bring adhesion weakening, which, together with elevated pulling, starts the sliding.

*CNO3 rescue of* *GEF-H1 knockdown*: We model this experiment by 1.5-fold increase of the basal myosin activation rate
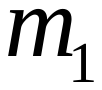
and by making the GEF-H1-dependent myosin activation rate,
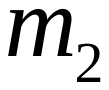
, equal to zero. The model predicts, correctly, that the FA sliding takes place (Supplemental Figure). The explanation is the same as in the previous case.

*Treatments by Y27632 or blebbistatin*: We model these experiments by 5-fold decrease of the basal myosin activation rate
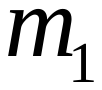
and by a significant decrease (Supplemental Table 2) of the GEF-H1-dependent myosin activation rate,
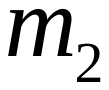
. The model predicts, correctly, that the FA sliding does not occur (Supplemental Figure). The explanation is straightforward: despite the force increase due to the GEF-H1-dependent myosin activation, even the maximal developed force is too weak to slide the adhesion even against the weakened adhesion.

Lastly, we have not simulated inhibition of FAK, Kinesin-1, or APC, but it is clear qualitatively that the model can predict no FA sliding in these cases assuming that these inhibitions abolish adhesion weakening by MTs. This can prevent the sliding despite the transient increase of the force or permanent force increase, as in the *CNO3* case.

**Table S1:** model variables

| 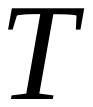 | Number of microtubule tips on the adhesion |
| --- | --- |
| 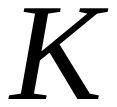 | Number of KANK molecules on the adhesion |
| 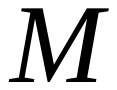 | Number of activated MLCs proximal to the adhesion |
| 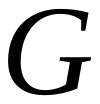 | Activated GEF concentration proximal to the adhesion |
| 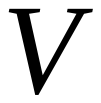 | Rate of slippage of the adhesion |
| 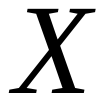 | Displacement of the adhesion |
| 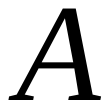 | Number of vinculin molecules in the adhesion |

**Table S2**: model parameters

| 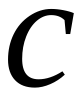 | Rate of microtubules’ leaving the adhesion | 1/30 sec |
| --- | --- | --- |
| 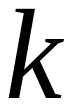 | Rate of KANK dissociation | 1/60 sec |
| 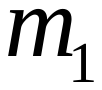 | Basal rate of MLC activation | 1 (control,GEFKD), 1.5 (CNO) 0.2 (BlebY) |
| 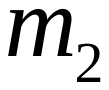 | GEF-H1-dependent rate of MLC activation | 5 (control), 0.2 (BlebY), 0.1 (CNO), 0 (GEFKD) |
| 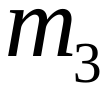 | Basal rate of MLC deactivation | 1/30 sec |
| 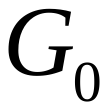 | Basal GEF-H1 concentration in the cytoplasm | 0.5 |
| 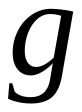 | Sequestered/released GEF-H1 amount per microtubule | 0.01 |
| 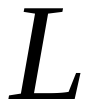 | Distance between the adhesion and myosin assembly site | 4 μm |
| 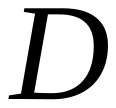 | GEF-H1 diffusion coefficient in the cytoplasm | 1 μm^2^/sec |
| 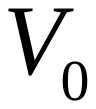 | Characteristic slippage velocity | 0.5 μm/sec |
| 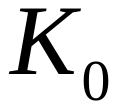 | Basal strength of a weak adhesion | 0.2 |
| 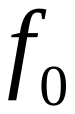 | Slippage force/adhesion ratio | 7.5 |
| 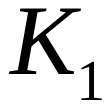 | Rate of KANK-adhesion association upon illumination | 2.2 |
| 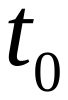 | Characteristic time of KANK loading upon illumination | 20 sec |
| 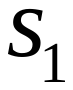 | Inverse threshold width for KANK loading function | 0.1 |
| 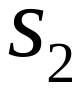 | Inverse threshold width for GEF-H1-dependent rate of MLC activation | 0.05 |
| 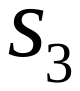 | Inverse threshold width for force-dependent rate of adhesion slippage | 1 |
